# ^89^Zr-oxine labelling and PET imaging shows lung delivery of a cell/gene cancer therapy

**DOI:** 10.1101/736967

**Authors:** P. Stephen Patrick, Krishna K. Kolluri, May Z. Thin, Adam Edwards, Elizabeth K. Sage, Tom Sanderson, Benjamin D. Weil, John C. Dickson, Mark F. Lythgoe, Mark Lowdell, Sam M. Janes, Tammy L. Kalber

## Abstract

**Purpose:** MSCTRAIL is a new stem cell-based therapy for lung cancer, currently in phase I evaluation (ClinicalTrials.gov ref: NCT03298763). Biodistribution of cell therapies is rarely assessed in clinical trials, despite cell delivery to the target site often being critical to presumed mechanism of action. This preclinical study demonstrates that MSCTRAIL biodistribution dynamics can be detected non-invasively using ^89^Zr-oxine labelling and PET imaging, thus supporting use of this cell tracking technology in phase II evaluation.

**Methods:** MSCTRAIL were radiolabelled with a range of ^89^Zr-oxine doses, and assayed for cell viability, phenotype and therapeutic efficacy post-labelling. Cell biodistribution was imaged in a mouse model of lung cancer using PET imaging and bioluminescence imaging (BLI) to confirm cell viability and location *in vivo* up to 1 week post-injection.

**Results:** MSCTRAIL retained therapeutic efficacy and MSC phenotype at doses up to and above those required for clinical imaging. The effect of ^89^Zr-oxine labelling on cell proliferation rate was dose and time-dependent. PET imaging showed delivery of MSCTRAIL to the lungs in a mouse model of lung cancer, with PET signal correlating with the presence of viable cells as assessed by bioluminescence imaging, *ex vivo* autoradiography and matched fluorescence imaging on lung tissue sections. Human dosimetry estimates were produced using simulations and preclinical biodistribution data.

**Conclusion:** ^89^Zr-oxine labelling and PET imaging present an attractive method of evaluating the biodistribution of new cell-therapies, such as MSCTRAIL. This offers to improve understanding of mechanism of action, migration dynamics and interpatient variability of MSCTRAIL and other cell-based therapies.

## Introduction

Lung cancer is the leading cause of cancer death worldwide, survival rates are among the lowest [1], and improvement of treatment options is among the slowest of the major cancer types. The need for rapid translation and validation of new lung cancer therapies is therefore of high importance. Cell-based therapies have the potential to answer unmet clinical needs in a number of disease areas including oncology [2]. However the spatial and temporal distribution of transplanted cells is rarely assessed in patients due to a lack of established technologies [3]. This can lead to concerns over safety and efficacy, and delays in translation [3-5]. We assess here the suitability of ^89^Zr-oxine labelling and PET imaging to track MSCs (mesenchymal stromal cells) in a clinical cell/gene-therapy trial for lung cancer.

TACTICAL (Targeted stem cells expressing TRAIL as therapy for lung cancer) is a prospective, randomised phase I/II trial to assess the safety and efficacy of third party, pooled allogeneic MSCs expressing TRAIL (MSCTRAIL) in combination with Pembrolizumab, Cisplatin and Pemetrexed as first line therapy for metastatic lung adenocarcinoma [6, 7] (ClinicalTrials.gov Identifier: NCT03298763). TRAIL (TNF-Related Apoptosis-Inducing Ligand) selectively induces apoptosis in cancer cells via binding to cell surface death receptors, thereby activating the extrinsic apoptotic pathway [8]. The clinical utility of soluble recombinant TRAIL is limited by its short half-life in the body. This can be overcome by MSCs constitutively expressing TRAIL in the tumour microenvironment and their tumour tropism [8-10]. This study is a preclinical assessment of the feasibility of imaging lung-specific MSCTRAIL delivery, and duration of cell retention, using ^89^Zr-oxine and PET in a separate imaging arm of the TACTICAL phase II trial.

^89^Zr-oxine has recently emerged as a favourable PET alternative to ^111^In-oxine [11-13], which has enabled diagnostic tracking of white blood cell infusions for over 40 years with SPECT or scintigraphy. In addition to offering ^111^In-oxine’s advantages of ∼3 day half-life and rapid radiolabelling, ^89^Zr-oxine benefits from the >10-fold increased sensitivity of detection associated with PET [14]. With the introduction of total-body clinical PET scanners this is set to increase by a further ∼40-fold, reducing scan times and radioactive doses for patients and thus increasing the practicality of whole-body cell tracking in the clinic [15]. A handful of preclinical studies have so far shown the worth of ^89^Zr-oxine in tracking cell therapies in mouse models, including T-cells [16, 17], dendritic cells, NK cells [13], and bone marrow cells [18, 19]. However ^89^Zr-oxine toxicity is dose-dependent and varies between cell types, requiring individual evaluation with each prospective cell therapy prior to clinical implementation.

Therapeutic and phenotypic equivalence after ^89^Zr-oxine labelling was shown for MSCTRAIL, and dose-dependent toxicity investigated to ascertain tolerated labelling doses. Delivery of viable cells and their retention in the lung was shown over 7 days in a preclinical lung tumour model, and human dosimetry estimates were calculated to facilitate translation. This study illustrates the feedback that ^89^Zr-oxine labelling and PET imaging could provide on the bio-distribution of cell-based therapies during the clinical trial phase of development. Wider use of cell tracking techniques such as this promise to contribute to a better scientific understanding of therapeutic cell behaviour within patients, as well as their associated safety profile, thereby informing future developments and effective translation.

## Results

### ^89^Zr-oxine effect on viability of MSCTRAIL is dose and time dependent

Umbilical cord tissue-derived mesenchymal stromal cells (uct-MSCs) were transduced using a lentiviral vector encoding TRAIL. TRAIL expression at 95% was confirmed by flow cytometry (Figure S1), and these cells (hereafter referred to as MSCTRAIL), were used for all subsequent experiments except where otherwise stated.

To assess ^89^Zr-oxine cytotoxicity, freshly harvested MSCTRAIL were radio-labelled over a 10-fold range of doses between 1515 and 152 kBq/10^6^ cells (as measured after 3 washes), or sham labelled in PBS alone, or PBS + 3% DMSO (used as a ^89^Zr-oxine vehicle). A 20 minute labelling incubation was chosen to facilitate use within the 90 minutes post-defrosting during which TRAIL-MSCs typically maintain maximum viability when left in the cryopreservant [20]. Labelling efficiency correlated negatively with ^89^Zr-oxine dose (R^2^ =0.804), with the lowest two doses achieving the highest labelling efficiency with between 29 and 33% retained after 3 washes (Figure S2A), comparable to prior reports using a 30 minute incubation [12]. The highest dose showed the earliest effect on viability vs unlabelled cells after 3 days, though all doses resulted in a significant reduction in cell growth as measured by ATP and NADH metabolism from day 4 onwards (Figure S2B, C). Two-Way ANOVA analysis showed significant individual as well as interactive effects from time and dose (p<0.001).

Cryopreserved MSCTRAIL doses are thawed immediately prior to patient transfusion in the TACTICAL trial [20], and will be radio-labelled between thawing and transfusion for the imaging cohort. For ^89^ Zr doses equivalent to 37 to 100 MBq per patient receiving a cell dose of 5×10^6^ cells/kg (4×10^8^ cells for a patient of 80 Kg), we assessed the effects of labelling MSCTRAIL doses immediately after defrosting encompassing the clinical range of 92.5-250 kBq/10^6^ cells. The highest dose (332 kBq/10^6^ cells) showed the earliest effect on ATP metabolism vs sham (PBS)-labelled cells at 4 days post labelling, though all doses resulted in a significant reduction in ATP and NADH levels from day 7 (Figure 1 A,B). Time accounted for 74 % (ATP) and 68 % (NADH) of variation, dose for 9 % (ATP) and 13 % (NADH) of variation, with significant interaction between dose and time accounting for 11 % (ATP) and 17 % (NADH) of variation (2-way ANOVA, p<0.0001).

**Figure 1.**
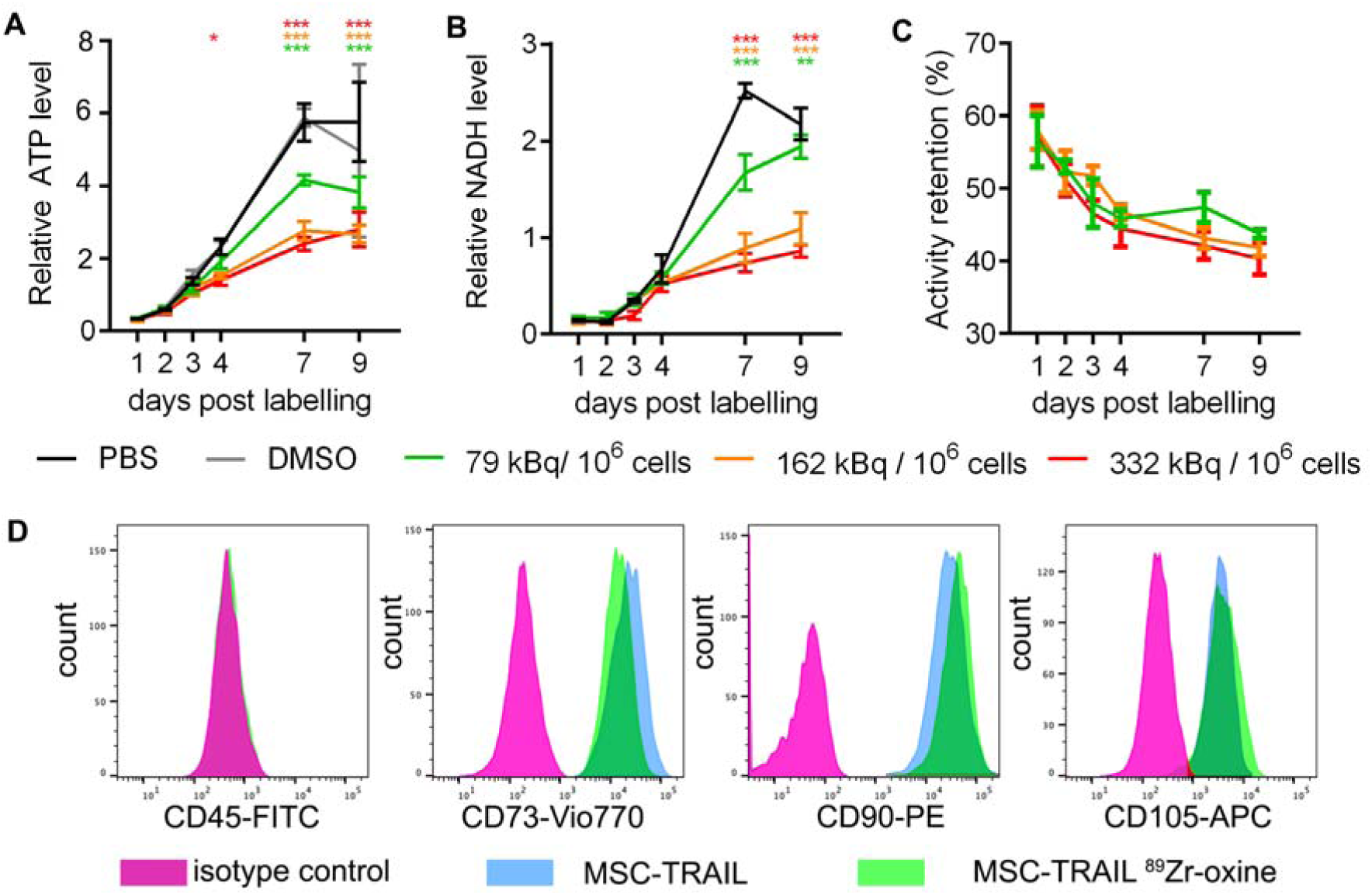
MSCTRAIL show time and dose-dependent sensitivity to ^89^ Zr-oxine labelling but retain MSC phenotype at high doses. MSCTRAIL cells were labelled from frozen with ^89^ Zr-oxine doses between 79 kBq/10^6^ and 332 kBq/10^6^. Cells show reduced proliferation with increasing dose as indicated by metabolism of ATP **A**. and NADH **B**. Error bars show standard deviation (SD). *p<0.05, **p<0.01, ***p<0.001 2-Way ANOVA with Dunnett’s multiple comparisons test vs PBS control. **C**. Retention of ^89^ Zr oxine decreases over time (n=4). 2-way ANOVA analysis showed that the majority of variation (67.5%) was due to time (p<0.0001), with dose having a significant (p=0.015) but small effect (3% of variation) on activity retention. **D**. MSCTRAIL retain their MSC phenotype (CD45-ve, CD73, CD90, and CD105+ve), post radiolabelling with ^89^Zr-oxine (dose shown 332 kBq/10^6^ cells).

The effect of dose on activity retention was also investigated (Figure 1 C), with the majority of variation in retention (67%) due to time (2-way ANOVA; p<0.0001), with a small but significant effect from dose (3%, p=0.015). Most of the label loss occurred during the first 24 hours after which label loss was slower, with only a further 12% being lost between 24 hours and 7 days.

Similar interactive and individual time and dose-dependent effects on ATP and NADH metabolism were observed in MSCTRAIL labelled directly after harvesting from culture (Figure S3A,B) across a comparable range of ^89^Zr-oxine doses (72 to 283 kBq/10^6^ cells). Labelling efficiency of frozen MSCTRAIL at 37.7 % (SD=2.2) was comparable to the mean of 43 % (SD=3.6) achieved with cells harvested directly from culture.

Since radiolabelling affects MSCTRAIL cell viability at higher doses, we investigated if this was due to cytotoxic or cytostatic effect induced by the radiolabel. Cell cycle analysis of MSCTRAIL labelled directly after thawing (Figure S4) showed changes in cell cycle profile from the 3 day to 7 day post labelling time points which increased with dose, consistent with metabolic changes (figure 1). The proportion of cells in G2/M in the top two radiolabelling doses doubled compared to the PBS control, suggesting that activation of the DNA damage checkpoint may be responsible for the decreased rate of proliferation at these doses. At the lower dose (79 kBq/10^6^ cells), cells showed a similar cell cycle profile to control cells at day 3 and 7, consistent with their more similar metabolic profile to unlabelled cells. Increases in apoptotic cell fraction (up to 32% for the higher doses) were only seen at the highest two doses at day 7, though ∼7% of cells were still in S phase, consistent with the slow but continuing replication up to day 7 (Figure 1).

To further assess the effects of radiolabelling, Western blot analysis of cell extracts from 7 days post-labelling were assessed for γ-H2AX upregulation for DNA damage signalling, and Nrf2 upregulation for oxidative stress (Figure S5A). Protein upregulation was not detected in radiolabelled compared to control cells at this time point, suggesting that DNA damage has either been repaired in cells by this time point or that it is occurring below the threshold of detection, and that cells are not showing detectable levels of oxidative stress signalling. To confirm the functioning of these homeostatic signalling pathways in cells after radiolabelling, this experiment was repeated using TBHP to induce DNA damage and oxidative stress, which showed upregulation of both proteins in radiolabelled and non-radiolabelled cell populations (Figure S5B).

### Radiolabelling does not affect MSC-specific cell surface marker profile

Flow cytometry was used to evaluate the effect of radiolabelling on MSCTRAIL surface-marker phenotype, using the ISCT-approved MSC identification panel of antibodies [21]. Low to high dose radiolabelled and control (PBS and PBS+DMSO sham labelled) MSCTRAIL showed the expected cell surface marker expression for MSCs (+ve for CD73, CD90, and CD 105; -ve for CD14, CD19, HLA class II, CD34, and CD45), (Figure 1D and S6). Radiolabelling doses above and below the range needed for clinical PET imaging of MSCTRAIL are therefore compatible with maintaining an MSC-like cell-surface marker phenotype.

### TRAIL expression and therapeutic function is unaffected by radiolabelling

To assess the effect of ^89^ Zr-oxine on therapeutic capacity, maintained TRAIL expression on MSCs was confirmed 7 days post-labelling with Western Blot (Figure 2A). Labelled and unlabelled MSCTRAIL were co-cultured with luciferase-expressing TRAIL-sensitive (NCI-H28, human lung mesothelioma) and partially TRAIL-resistant (PC9; human lung adenocarcinoma) cancer cells (Figure 2B). Reduction in viable cancer cell population was measured as light output following the addition of luciferin, and compared to untreated control populations. Soluble recombinant TRAIL at 10 and 50 ng/mL was used as a positive control. Population reduction of viable cancer cells was further confirmed by light microscopy (Figure S7A). Radiolabelled MSCTRAIL maintained apoptosis-inducing ability against both cancer cells lines (figure 2B). An orthogonal cell death assay by Annexin V/DAPI flow cytometry in four cancer cell lines (H28, PC9, MDAMB-231, H2869) also confirmed maintained induction of apoptosis by MSCTRAIL post radiolabelling, giving equivalent results (Figure 2 C and D, S7B).

**Figure 2.**
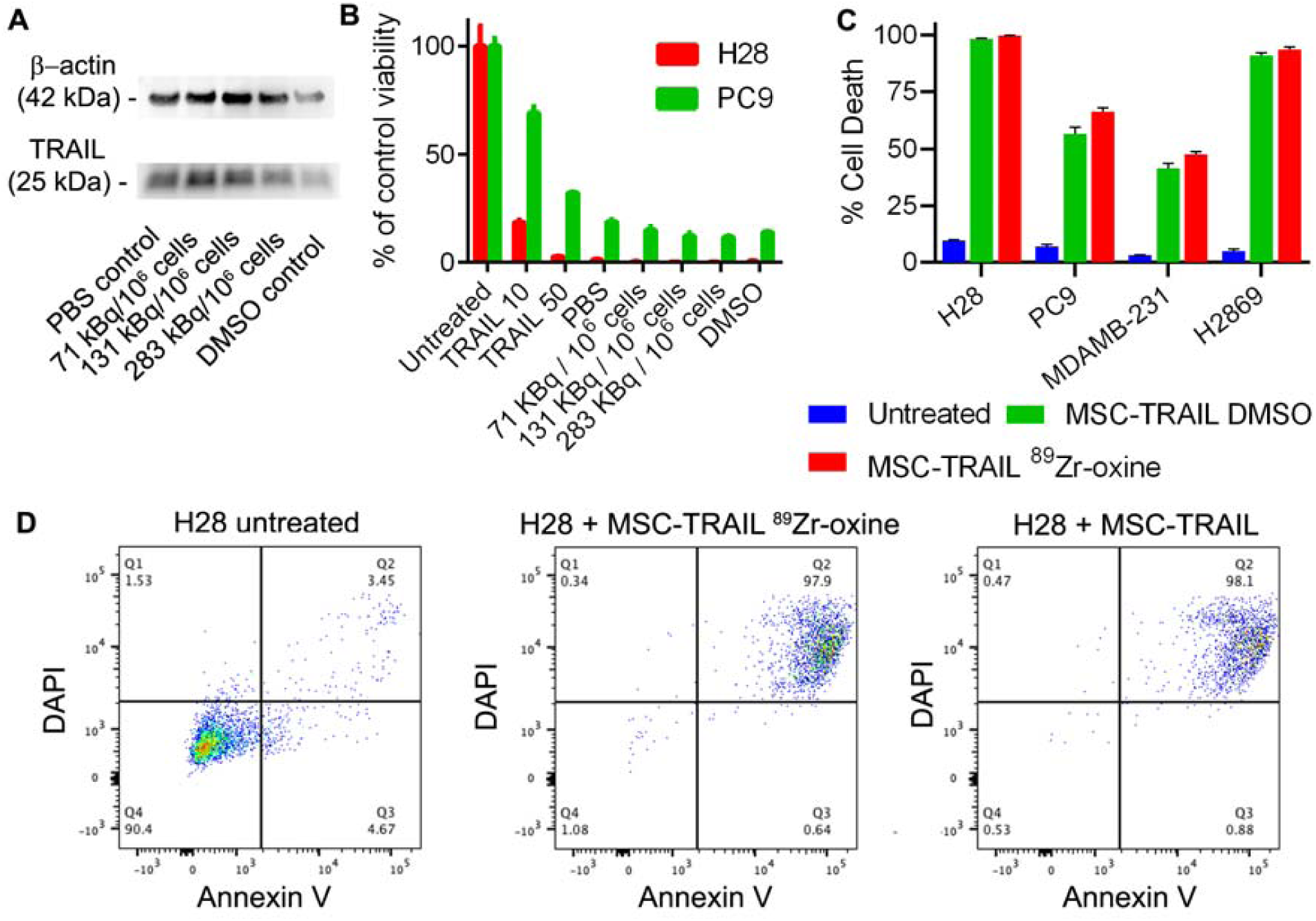
TRAIL expression and therapeutic function is maintained following radiolabelling. **A**. Western blot analysis on protein extracts 7 days post-radiolabelling shows maintained TRAIL expression. **B.** Viability of two human lung cancer cell lines (NHI-H28 and PC9) were measured using luciferase bioluminescence after treatment with soluble trail at 10 or 50 ng/mL, or co-incubation for 24 hours with MSCTRAIL labelled 3 days prior with the indicated doses. Bars show the mean of 4 populations, error bars show SD. **C**. Apoptosis of four cancer cell lines untreated or after incubation with MSCTRAIL or MSCTRAIL radiolabelled with ^89^ Zr-oxine at 332 kBq/10^6^ cells, measured using Annexin V staining and flow cytometry. Error bars show SD, n=3. **D**. Representative FACs plots from C, showing apoptosis of H28 cells following incubation with control or radiolabelled MSCTRAIL.

### *In vivo* TRAIL-MSC tracking in a mouse model of lung mesothelioma

Immunocompromised mice were implanted intra-pleurally in the right hand side with a human mesothelioma cell line (luciferase-transduced CRL-2081), and tumour growth was followed using bioluminescence imaging (supplementary figure 7). T_2_-weighted magnetic resonance imaging and CT showed localisation of the tumour to the right lung (figure 3 A to C) at 15 days post-implantation. MSCTRAIL were then thawed, radiolabelled (311 kBq/10^6^ cells), and injected intravenously (1.5×10^6^ cells). PET-CT showed ^89^Zr signal throughout the lung, including within the area containing the tumour, (figure 3 B to D).

**Figure 3.**
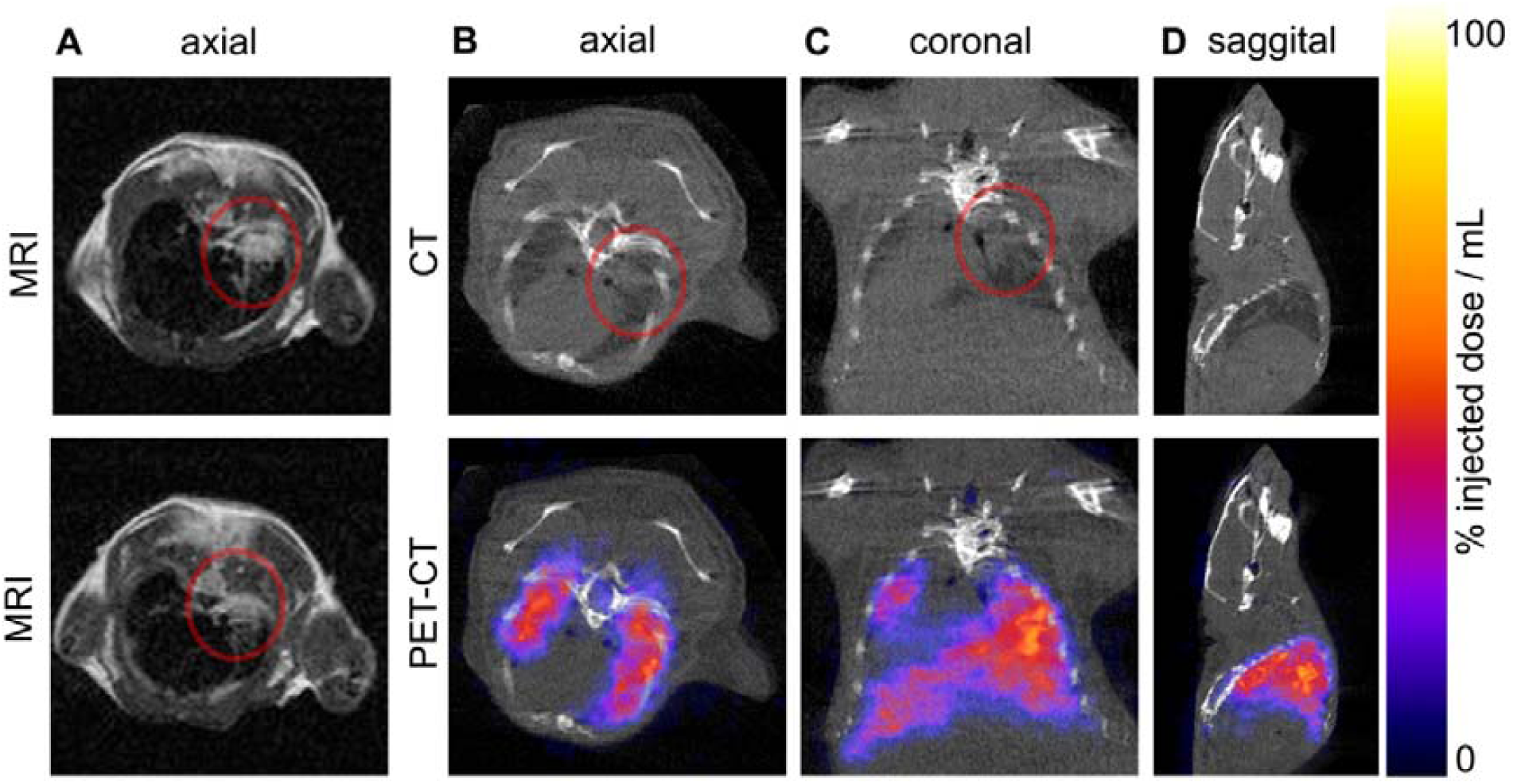
Tumour visualisation with MRI and CT, and location of ^89^Zr-oxine labelled MSCTRAIL visualised with PET at 2 days post transplantation. Lung tumours detected with **A.** axial T_2_-weighted magnetic resonance imaging (T_2_ RARE TE=55ms, respiratory gated) showing two consecutive 1 mm thick slices through the lung tumour (shown in the red circle) at 15 days post implantation **B**. axial, **C**. coronal, and **D**. sagittal CT slices, with corresponding PET overlay showing the location of ^89^Zr-oxine labelled MSCTRAIL in the lungs and liver.

PET-CT at 1 hour, 1, 2, and 7 days post-injection enabled visualisation and quantification of MSCTRAIL biodistribution dynamics (figure 4). The majority of signal (60%) was found in the lung at 1 hour before decreasing while liver signal increased. From 1-7 days post injection, the proportion of the ^89^ Zr signal in the lung fell further from 24.6% (+/-10.4 % SD) to 16.0 % (+/-7.9% SD). Uptake in the spleen was also visible and peaked at 2.6% of the total injected dose (+/-0.64% SD) at 48 hours.

**Figure 4.**
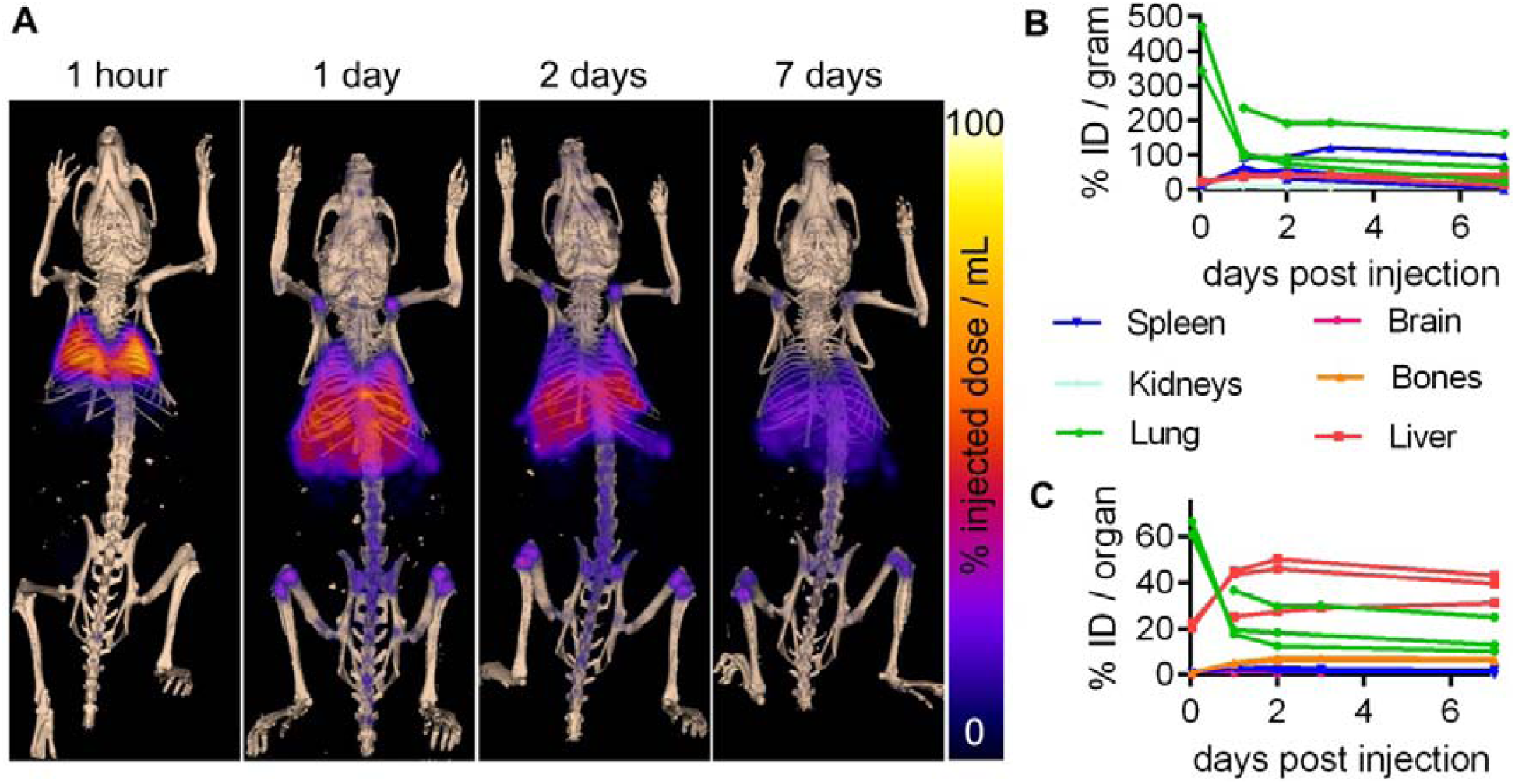
Whole-body biodistribution of ^89^ Zr-oxine labelled MSCTRAIL followed up to 1 week post-implantation. **A.** maximum intensity-projection PET showing ^89^ Zr-oxine-labelled MSCTRAIL overlaid on 3D-rendered bone CT at the indicated time points after intravenous injection. The indicated organs and bones were segmented using CT data and radioactivity quantified from PET data, giving **B**. % injected dose (ID) per gram, using the wet weight of tissue from each animal (n=3) following dissection at day 10 post MSCTRAIL injection, and **C**. % ID per organ.

**Figure 5.**
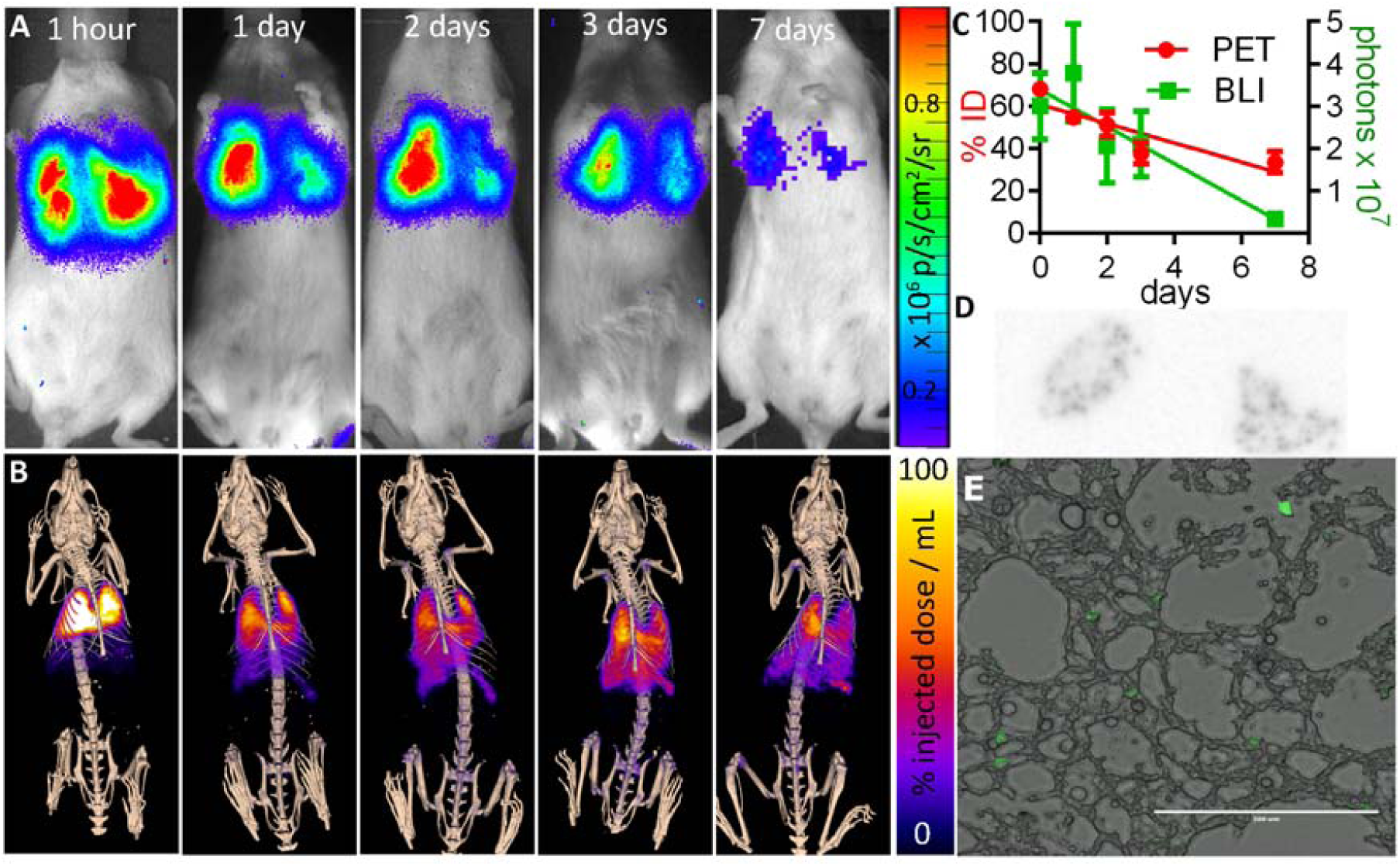
Viable MSC location correlates with PET signal. Intravenously injected ^89^Zr-oxine labelled uct-MSCs expressing luciferase and ZsGreen were tracked over 7 days using **A**. Bioluminescence imaging, and **B**. PET-CT imaging. **C**. ROI analysis of PET and BLI data showed a comparable decrease in %ID and photons in the lungs over time (n=3, error bars SD). At 7 days post injection, lungs were removed and cyrosectioned, before imaging with **D.** Autoradiography, and E. Fluorescence microscopy, to confirm the co-location of ZsGreen-expressing cells with ^89^Zr signal within the lungs.

At ten days post MSCTRAIL injection, organs were removed, weighed, and ^89^Zr activity was measured. Activity per organ was normalised to organ weight and decay-corrected (Figure S9 and table 1). Consistent with PET-CT and ROI analysis, high amounts of activity in the lung and liver were found, with the lung having the most activity per gram (78% ID per gram +/-26%) followed by the liver (32.8 % ID per gram +/-8.6 %), though the liver had a higher absolute percentage of the injected dose (29.4% +/-11.5%) compared to the lungs (12.4 % +/-4.3%). Activity uptake in the spleen was also still high at 10 days post injection (43.9 % +/-5.3 % per gram), consistent with its visibility in the PET-CT scans (figure 4A), however in absolute terms this represented just 1.6% +/-0.5% of the total injected activity.

### Correlation of PET signal with viable cell location

If ^89^Zr-oxine labelling is to be of practical use in tracking cell biodistribution, PET signal must correlate with viable cell location over biologically-relevant timeframes. To evaluate this, uct-MSCs were transduced to express luciferase and the green fluorescent protein ZsGreen to enable independent viability and/or location confirmation with *in vivo* bioluminescence imaging (BLI) and *ex vivo* fluorescence microscopy. MSCs expressing ZsGreen-luciferase were then radiolabelled with ^89^Zr-oxine (412 kBq/10^6^ cells), injected intravenously, and imaged with PET-CT and BLI for 7 days. This showed correlation of radiolabel with viable cell location, with an initial predominance of signal in the lung which decreased over time on PET and BLI (5A-C). Organs were then removed, cryosectioned, and imaged with fluorescence microscopy and autoradiography (5 D and E), together with DAPI staining (Figure S10), confirming the presence of the transplanted ZsGreen-expressing uct-MSCs in the lungs. Despite a significant fraction of injected radioactivity being measured in the liver and spleen at the end time point, no signal was detected in these organs with *ex vivo* bioluminescence (Figure S11A,B). However, examination of tissue sections with fluorescence microscopy did suggest the presence of debris from ZsGreen-expressing cells (S11D,E). This is consistent with the liver uptake of intravenously-injected heat-inactivated MSCs seen with PET-CT (S12).

### Human dosimetry estimates

To anticipate clinical use, human dosimetry estimates were calculated with OLINDA software [22] using mouse to human extrapolations according to Stabin [23] and the preclinical *in vivo* region of interest analysis data and *ex vivo* biodistribution data (see Table S2 to S4). For an injected dose of 37 MBq this gave mean effective dose estimates for male and female patients of 32.2 and 41.4 mSv, respectively. For 100 MBq per patient this corresponds to an effective dose of 87.1 and 111.8 mSv for male and female patients respectively. The organ specific dose is estimated to be highest in the lungs (5.09, 6.58 mSv/MBq), spleen (2.12, 2.57 mSv/MBq), and liver (1.86, 2.39 mSv/MBq) for male and female patients respectively.

## Discussion

Many factors potentially contribute to the complexity of cell behaviour and cell/host interactions including cell source and pre-processing, injection route, patient age, immune system, co-morbidities, genetics, life history, and microbiota [24-26]. Without assessing cell biodistribution in patients using cell tracking techniques, it remains difficult to evaluate the effect of these variables on cell behaviour, and the failure of many emerging cell therapies [27].

To support integration of ^89^Zr-oxine cell tracking into the TACTICAL trial we have shown that TRAIL-expressing umbilical cord tissue-derived MSCs (MSCTRAIL) can be tracked non-invasively to the lungs in a preclinical lung cancer model up to 7 days post-injection. PET signal correlated with viable cell signal from bioluminescence imaging, increasing confidence in the reliability of this technique. Labelling was achieved from frozen cell stocks with relevant doses of ^89^Zr-oxine within 45 minutes - below the 90 minutes over which frozen MSCTRAIL retain optimal viability post-thawing [20], suiting this approach to translation.

PET imaging of ^89^ Zr-antibodies has been reported up to 5 days post injection with 37 MBq/patient [28, 29], and up to 6 days using 70 to 75 MBq/patient [30, 31]. To achieve 37 MBq/patient receiving 5×10^6^ MSCTRAIL cells/kg (ie 4×10^8^ cells for an 80 Kg patient), this equates to ^89^Zr-oxine labelling at 92.5 kBq/10^6^ cells - towards the lower range of doses assessed here; for 70 MBq/patient this is 185 kBq/10^6^ cells. Even at higher doses TRAIL expression, therapeutic efficacy, and MSC cell-surface markers were retained. On the other hand, decreased proliferation rates were seen by day 4 to 7. Proliferation of MSCTRAIL during TACTICAL is not however expected due to their use in combination with Cisplatin and Pemetrexed.

Currently, bone marrow-derived MSCs (bm-MSCs) predominate among approved MSC-based cell therapy products, but their use in clinical trials relative to other MSC sources has more than halved since their peak [32, 33]. Conversely, the use of umbilical cord tissue-derived MSCs for cell therapy, as in TACTICAL, is increasing due to various advantages [32, 34, 35]. Sourcing of cord donor material is non-invasive, unlike harvesting bone marrow, while production scale-up for cell therapy is eased by their delayed senescence onset, lack of contact inhibition enabling higher harvest densities, and faster proliferation rate. Allogenic use of uct-MSCs is facilitated by their wide multi-potency and hypo-immunogenicity. Despite sharing a similar transcriptomic profile and beneficial phenotype with embryonic stem cells, uct-MSCs do not possess their ability to form teratomas.

Aside from the present study on umbilical cord tissue-derived MSCs, ^89^Zr-oxine-labelling and PET imaging has also been demonstrated with T-cells, dendritic cells, and bone marrow cells [12, 13, 17, 19]. Together, these studies illustrate the benefits of ^89^Zr-oxine labelling: fast implementation, non-invasive tracking over a week post-injection, and interpretation of images un-complicated by the degree of background signal given by the systemically-administered tracers used with nuclear reporter-gene systems [36]. Though a modest amount of radio-label is inevitably lost from cells, the bone-specific uptake of free ^89^Zr is well-characterised and reduces ambiguity during analysis [37]. However, several limitations to this imaging strategy also need to be considered. Firstly, the doses required for tracking cells may reduce proliferation, and this effect varies between cell-types, and time post-labelling. Secondly, though a good correlation between PET signal and viable cell location was shown here using bioluminescence imaging, ^89^Zr-oxine signal does not directly indicate cell viability or proliferation. Hence for rapidly and non-uniformly expanding cell populations, genetic labelling strategies may be more appropriate [36, 38].

## Conclusion

This study shows that umbilical cord tissue-derived MSCs, and MSCTRAIL derived from these, can be radiolabelled with ^89^ Zr-oxine. Phenotype and therapeutic effect post-labelling were retained, and cells could be tracked *in vivo* up to 7 days using PET. This supports the feasibility of a first-in-man ^89^ Zr-oxine cell-labelling arm in phase II of the TACTICAL clinical trial, where it should lead to a better understanding of this experimental cell therapy, its whole-body biodistribution dynamics, and inter-patient variability.

## Methods

### ^89^Zr-oxine synthesis and purification

^89^Zr-oxalate stock (2-40 MBq; Perkin Elmer) was diluted to 500 μL with HPLC-grade water and neutralised with 1M NaOH (Sigma-Aldrich). To this was added 20 μL of a 10 mg/mL solution of 8-hydroxyquinoline (ACS reagent, 99%, Sigma-Aldrich: 252565) in chloroform (Sigma-Aldrich), in a screw-top round-bottomed tube (BRAND® culture tubes, 6.5 mL, AR-Glas®, Sigma-Aldrich) held onto a vortex mixer (Grant Bio, model PV-1) for 5 minutes using a clamp. Chloroform was added (480 μL) before vortexing for 25 minutes. After brief centrifugation the chloroform phase containing Zr-oxine was removed and evaporated at 80 ° C in conical bottom HPLC vial (Supelco CD vial, 9 mm screw, Sigma Aldrich), before resuspension in 15 μL of dimethyl sulfoxide (DMSO anhydrous, Sigma-Aldrich) at 50 °C for 20 minutes. Radiochemical yield (RCY) was calculated as the percentage of total original activity extracted into the chloroform phase, giving an average RCY of 74.2% ± 4.3 SEM (n=16). Negative control synthesis was performed without addition of 8-hydroxyquinoline to the chloroform phase, resulting in retention of all the activity in the aqueous phase.

### MSC isolation

Umbilical cord tissue was manually dissected, then enzymatically and mechanically digested to isolate MSCs using plastic adherence to cell culture-treated flasks. Passage-0 cultures were cultured in serum-containing α-MEM (Gibco) with antibiotic/antimycotic, then expanded subsequently in serum-free, xeno-free medium prior to labelling.

### Cell culture

Cells were cultured in 5% CO_2_, at 37 ° C, in either RPMI (Gibco), supplemented with 10% fetal bovine serum (FBS) for PC9 (lung cancer) CRL2081, H28, H2869, (malignant pleural mesothelioma) and MDAMB-231 (breast cancer) cells, or α-MEM (Gibco) with 10% FBS for uct-MSCs (MSCTRAIL). α-MEM was also used for MSC/cancer cell co-culture apoptosis assays. All tissue culture reagents were obtained from Invitrogen. For cell culture following radiolabelling, growth media was supplemented with 50 U/mL penicillin and 50 mg/mL streptomycin (Invitrogen).

### Cell labelling

MSCTRAIL (1.5×10^6^ per condition for dose toxicity experiments; 2×10^7^ for *in vivo injections*) were re-suspended in 200 μL PBS. A further 100 μL was added containing PBS + ^89^Zr-oxine re-suspended in DMSO (3% final concentration per 300 μL in both ^89^Zr-oxine and sham labelling conditions), with a further control of PBS alone. Cell suspensions were incubated for 20 minutes at room temperature, pelleted at 900 g, supernatant removed, and resuspended in PBS. This was repeated three times to remove unbound radiolabel. Final bound dose was measured using an ionization chamber (Curiementor 4, PTW Freiburg GmbH), and expressed as kBq per 10^6^ cells, following cell counting. Total labelling and washing time was ∼45 minutes on each occasion.

### Lentiviral production and transduction

Please see supplementary information.

### Flow cytometry

For flow cytometry detection of TRAIL expression, cells were stained with a 1:50 dilution of phycoerythrin (PE)-conjugated mouse monoclonal antibody against human TRAIL (550516, BD Biosciences). For immunophenotyping of MSCs the cells were stained with 1:50 dilution of fluorochrome-conjugated mouse monoclonal antibodies against CD73 (PE-Vio770, 130-104-224, Miltenyi Biotec), CD105 (APC, 130-094-926, Miltenyi Biotec), CD90 (PE, 561970, BD Biosciences), CD34 (FITC, 560942, BD Biosciences), CD45 (FITC, 560976, BD Biosciences) and HLA-DR (FITC, 555560, BD Biosciences).

### *In vitro* cell viability assays

MSCTRAIL were labelled as above with a range of doses of ^89^ Zr-oxine dissolved in DMSO, or DMSO only or PBS only for the control conditions. Cells were seeded into 96 well culture plates at 10,000 / well in 100 μL culture medium straight after radiolabelling, n=4 wells per condition. A separate plate was seeded for each time point and assay. For ATP measurement, CellTiter-Glo® reagent was freshly-prepared, and added at a 1:1 ratio to culture medium. Light output was measured using a plate reader (VarioSkan Lux, Thermofisher Scientific) 10 minutes after reagent addition. For measurement of the reducing power of cells (NADH levels), XTT reagent (Roche) was prepared according to manufacturer’s instructions. To each well 50 μL reagent was added, including control wells containing media but no cells, followed by 4 hour incubation at standard culture conditions. Absorbance at 490 nm and 630 nm was recorded using a plate reader (VarioSkan Lux, Thermofisher Scientific), and mean absorbance of each well at 630 was subtracted from absorbance at 490 nm, and corrected for background metabolism in control wells. For measuring the cell viability of cancer cells upon coculture with MSCTRAIL, ZS-Green luciferase expressing H28 and PC9 tumour cells were seeded into 96 well culture plates at 5000 cells/well in 100 μL culture medium, and treated with 2000 MSCTRAIL for 24 hours. To measure cell viability luciferin was added to each well (10 µL, 15 mg/mL) and photon counts were obtained after 10 minutes using a plate reader (VarioSkan Lux, Thermofisher), with 1 second acquisition per well. Each condition was obtained in triplicate.

### Activity retention assay

MSCTRAIL were seeded at 10,000/well in α-MEM (Gibco) with 10% FBS in 96 well plates. At each time point growth media was removed and pooled with a wash of 100 μL PBS (+ 1mM EDTA). Cells were lysed using 100 μL RIPA buffer (Thermo Scientific), which was pooled with a further wash of 100 μL PBS (+ 1mM EDTA). Activity in media and cell fractions was measured using a Wizard 2480 automated gamma counter (PerkinElmer), and the % activity retained calculated as the percentage of the total amount in the cell+media fractions in the cell fraction. Four individually seeded replicates were taken per time point.

### *In vitro* apoptosis assay

To assess the apoptosis of tumour cells co-cultured with MSCTRAIL, DiI-labeled cancer cells were plated into a 96-well plate (5000 cells/well), to which 2000 MSCTRAIL cells were added for 24 hours. Floating and adherent cells were stained with AF647-conjugated Annexin V (Invitrogen) and 2 μg/mL DAPI (Sigma) and were assessed by means of flow cytometry. Annexin V+ cells were considered to have undergone apoptosis; Annexin V+/DAPI+ cells were considered to be dead by apoptosis.

### Western Blot

Please see supplementary information.

### Animal work

Female mice with severe combined immunodeficiency (NOD-SCID Gamma, strain NOD.Cg-Prkdc scid Il2rgtm1Wjl/SzJ; Charles River, UK) were aged 6-8 weeks and weighed 20-22 g at the time of implantation. Procedures were carried out under the authority of project and personal licenses issued by the Home Office, UK, and were approved by local Animal Welfare and Ethical Review Bodies.

### Cell implantation

A small patch of fur was shaved over the right hand side of the rib cage. Under isoflurane anaesthesia 1×10^5^ Human mesothelioma cells (CRL2081, transduced to express luciferase) were injected into the intra-pleural space between the third and fourth rib up from the bottom of the ribcage in 50 μL PBS. Tumour growth was followed by bioluminescence imaging at 1, 7, 10, 16, and 22 days post injection, at which point mice were injected intravenously with 1.5 ×10^6 89^ Zr-oxine labelled MSCTRAIL in 200 μL PBS.

### Bioluminescence Imaging

Mice were anaesthetised with isofluorane and kept at 37 ° C, injected intraperitoneally with 150 mg/kg of D-Luciferin solution and imaged at 20 minutes post injection (for tumour monitoring) and 15 minutes post injection (for MSCs) using an IVIS Lumina (Perkin Elmer). Exposure times were optimised to ensure sufficient signal was obtained without saturation. Light output was quantified using ROI analysis and normalised to photons / second/ steradian. Cherenkov luminescence from ^89^Zr decay could not be seen above background noise, but was compensated for using images taken prior to luciferin injection with background signal subtracted over the relevant ROI from post-luciferin values.

### PET imaging

Mice were imaged at the stated time points after intravenous injected of ^89^ Zr-oxine labelled MSCTRAIL, using a PET-CT (Mediso NanoScan) interfaced to InterView Fusion software. CT was acquired at 50 kVp with 300 ms exposure, and reconstructed in 0.13 mm isotropic voxels. PET data was reconstructed in 5:1 mode using the Tera-Tomo algorithm in 0.4 mm isotropic voxels, and analysed using VivoQuant software (InViCro). 3D ROIs were drawn manually around the lungs, liver, kidneys, spleen, and brain, based on CT soft-tissue contrast, and bones were segmented using CT signal thresholding. The percentage of injected dose/organ (%ID/organ) was calculated using decay-corrected ROI values.

### Dosimetry estimation

Details of dosimetry calculations can be found in the supplementary information.

## Acknowledgements

P. S. P. acknowledges funding from the UK Regenerative Medicine Platform (MRC: MR/K026739/1) and MRC grant MR/R026416/1. T. L. K. is funded by an EPSRC Early Career Fellowship (EP/L006472/1). The TACTICAL trial is supported by MRC DPFS scheme grant MR/M015831/1. SMJ is a Welcome Trust Senior Fellow in Clinical Science (grant WT107963AIA) and is supported by the Rosetrees Trust, the Welton Foundation, the Roy Castle Lung Cancer Foundation and University College London Hospital (UCLH) Charitable foundation.

## Compliance with Ethical Standards

All applicable international, national, and/or institutional guidelines for the care and use of animals were followed. This article does not contain any studies with human participants performed by any of the authors. The authors have no conflicts of interest to declare.

## Supplementary Methods

### Western Blot

Cells labelled as above were seeded at a density of 7.5 million into T175 flask (Nunc) and split 1 in 5 at 72 hours, with the remaining cells being pelleted and snap frozen using powdered dry ice, and the re-seeded cells being harvested at 7 days post labelling and frozen in the same way. Frozen samples were left to decay until no background radiation was detectable, and protein lysates prepared using cell lysis buffer (RIPA, Thermofisher Scientific) according to the manufacturer’s instructions. Protein content of cell lysates were measured using the BCA assay according to the manufacturer’s instructions (Pierce, Thermofisher Scientific). Lysates were denatured using 10x reduction buffer (NuPage Sample reducing agent, Life Technologies), and loaded onto gels (BioRad Mini Protean TGX) at 30 mg together with 4x loading buffer (NuPage LDS sample buffer; Life Technologies), and pre-stained protein ladder (PageRuler™ 10 to 180 kDa; Thermofisher Scientific). Gels were blotted onto nitrocellulose membranes using the Trans-Blot Turbo device (BioRad), and probed using 1 in 1000 dilution of cleaved caspase 3 (Asp175) rabbit mAb antibody (Cell Signalling Technology, #9664) with horse-radish peroxidase conjugated anti-rabbit secondary antibody (Cell Signalling Technology, #7074), using the iBind Flex incubation system (Thermofisher Scientific). ECL developer (ECL Prime, GE) was added to membranes for 5 minutes, and carefully blotted off before imaging (ImageQuant, LAS4000; GE). Band intensity was measured using ImageJ (NIH.gov) with the gel analyse plug-in.

### Cell Cycle Analysis

Populations of MSCTRAIL labelled from frozen were analysed at 72 hours and 7 days post radiolabelling using a Chemometec NC-3000 benchtop image cytometer, using the DAPI-based cell cycle assay according to the manufacturer’s instructions.

### Lentiviral production and transduction

The lentiviral plasmid expressing TRAIL, pCCL-CMV-flT, previously described [39] was used to over-express TRAIL on MSCs. The ZsGreen-luciferase plasmid, pHIV-Luc-ZsGreen (a gift from Bryan Welm, Addgene plasmid #39196) was used for generating ZsGreen luciferase-expressing lentivirus. Lentiviral vectors were produced by co-transfection of 293T cells with construct plasmids together with the packaging plasmids pCMV-dR8.74 and pMD2.G using DNA transfection reagent jetPEI (Source Bioscience UK Ltd). Lentiviruses were concentrated by ultracentrifugation at 17,000 rpm (SW28 rotor, Optima LE80K Ultracentrifuge, Beckman Coulter, Brea, CA) for 2 hr at 4°C. Titres were determined via transduction of 293T cells with serial dilutions of virus, and 8 μg/ml Polybrene (Sigma-Aldrich). TRAIL and ZsGreen expression were assessed by flow cytometry. MSCs were transduced at a range of MOIs using 8 μg/ml Polybrene and transduction efficacy assessed by flow cytometry.

### Magnetic Resonance Imaging

Images were obtained using a 1T MRI system (ICON; Bruker BioSciences Corporation, Ettlingen, Germany), interfaced to a console running Paravision 5 software (Bruker). A 301mm mouse body solenoid RF coil (Bruker) was operated on transmit/receive mode. A rectal thermometer and respiratory pad provided physiological monitoring (SA Instruments, New York, USA), with temperature maintained via a water-heated bed. Multi-slice images of the lungs were acquired using a T_2_-RARE sequence, with effective TE of 55.8 ms (Echo train length =6), TR=1463 ms, 8 averages, 12 slices, field of view of 2.56×2.56 cm, 128×128 matrix, 1 mm slice thickness, 0.1 mm slice spacing, and respiratory gating. Total scan time was 4 minutes 5 seconds.

## Supplementary figures

**Figure S1.**
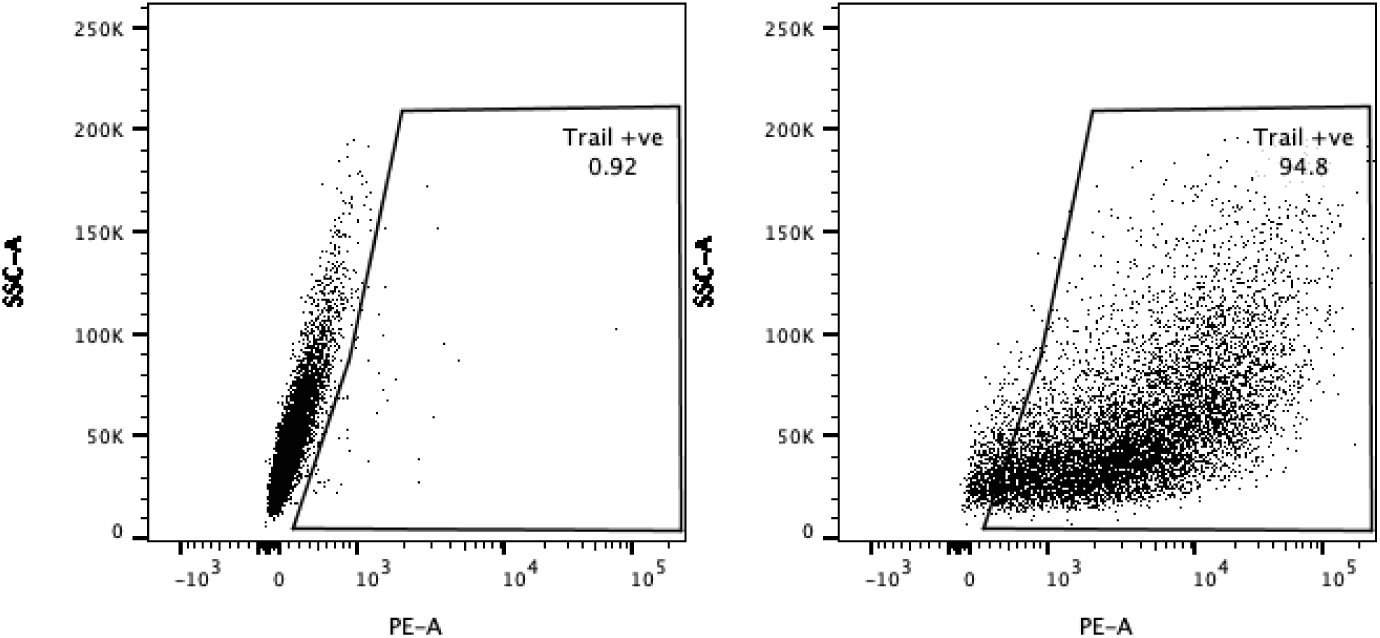
Flowcytometry analysis of. TRAIL expression of lentivirally-transduced cord-derived MSCs, showing 94.8% positive expression of TRAIL.

**Supplementary figure 2.**
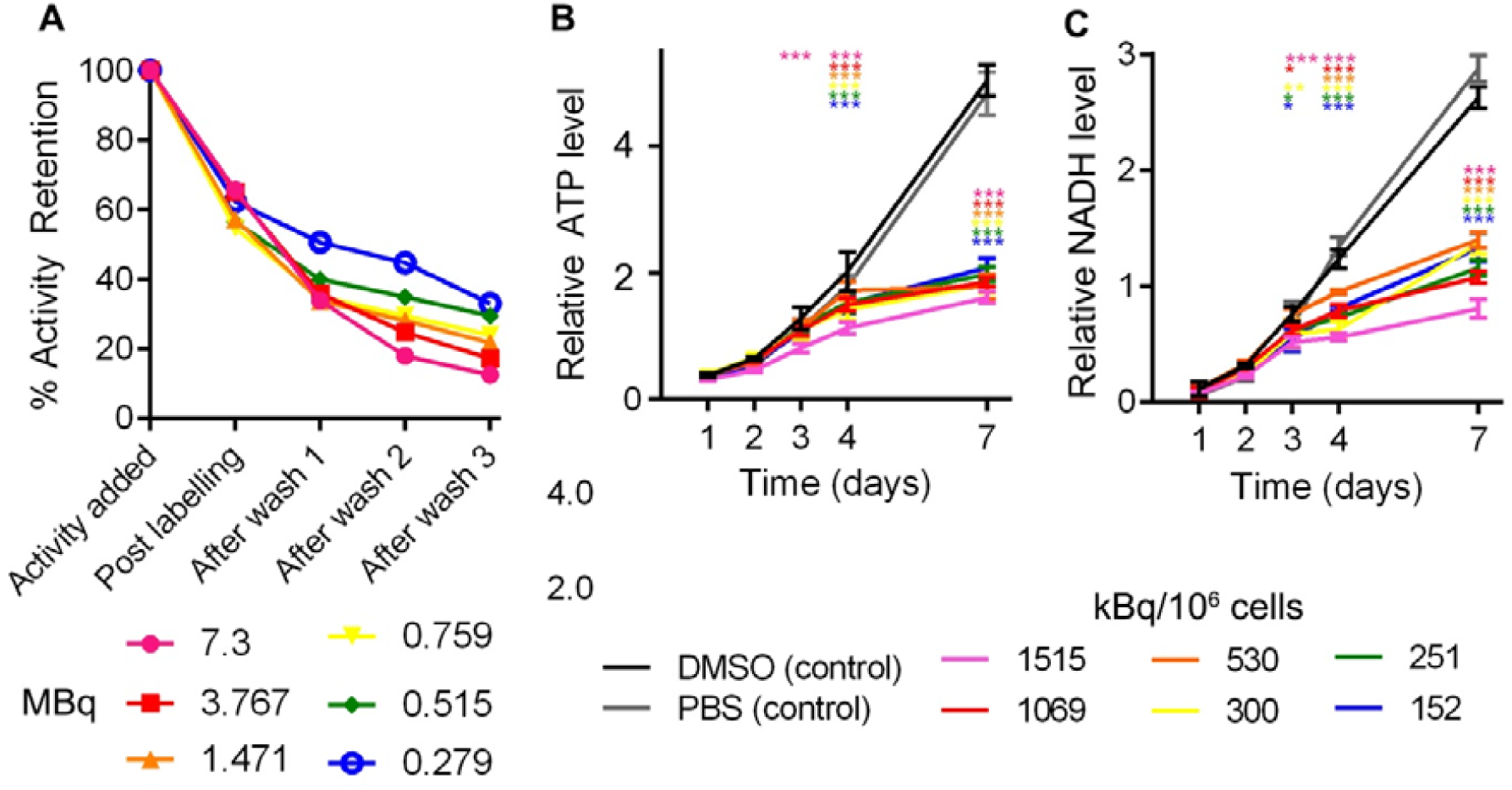
**(A)** 1.5 × 10^6^ cells were labelled in a final volume of 300uL, containing the indicated initial amounts of ^89^Zr-oxine (7.3 MBq to 0.279 MBq) and 3% DMSO. Three washes were found to be sufficient to wash the majority of the unbound activity from the cells. Final Labelling efficiency correlates negatively with the initial amount of ^89^Zr-oxine added to cells (R^2^=0.804). Effect of radiolabelling on cell metabolism was measured using assays for ATP (B) and NADH (C). Metabolism of labelled cell populations began to diverge from sham labelled (DMSO and PBS controls) populations at 3 days post labelling. Results were analysed using a 2-Way ANOVA with Dunnett’s multiple comparisons test comparing each population at a given time point against the PBS sham-labelled population * p<0.05 ** p<0.01*** p<0.001. No difference between the PBS and DMSO sham labelled populations was found at any time point. Dosing accounted for 10.4% (ATP) and 10.5% (NADH) of variation (p<0.001), Time for 62.6 (ATP) 69.8 (XTT) (p<0.001), with a significant interaction between time and dose accounting for 25.8% (ATP) and 18.8% (NADH) of variance (p<0.001). Points show the mean of 3 individually seeded populations, error bars show standard deviation (SD).

**Supplementary Figure 3.**
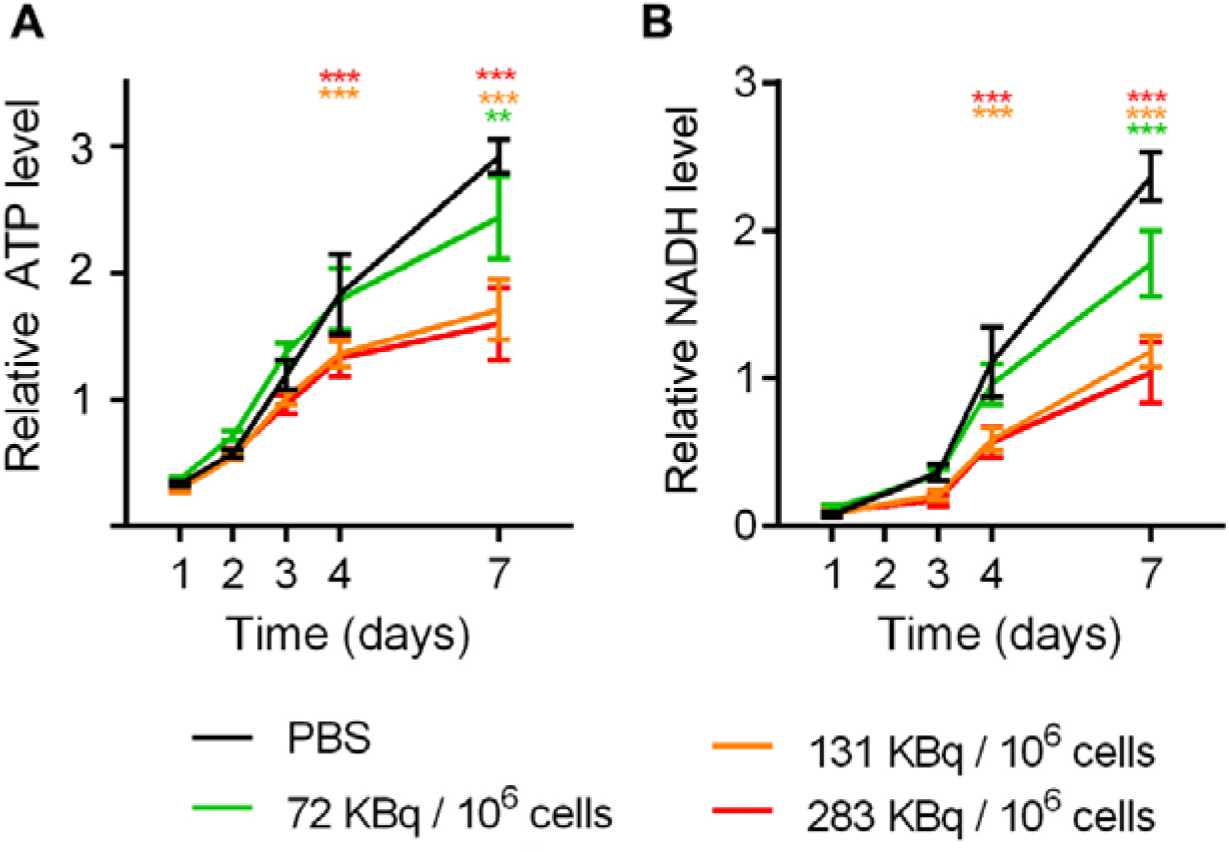
MSCTRAIL show time and dose-dependent sensitivity to ^89^ Zr-oxine labelling. MSCTRAIL were labelled following harvesting from culture with doses between 283 kBq/10^6^ cells and 72 kBq/10^6^ cells. Cells show reduced proliferation with increasing dose as indicated by metabolism of (A) ATP and (B) NADH. Dose, time, and dose-time interaction showed significant effect on variation (2-Way ANOVA; p<0.001). Time accounted for 74 % (ATP) and 83.7 % (NADH) of variation, dose for 9% (ATP) and 7% (NADH), with significant time/dose interaction of 5.6 % (ATP) and 7 % (NADH). Here the lowest dose (72 kBq/106 cells) showed a reduced effect on metabolism compared to the higher doses, with no significant difference in ATP or NADH metabolism compared to control cells until 7 days post labelling (p<0.01), whereas higher doses showed a difference by day 4 (p<0.01), Dunnett’s multiple comparisons test.

**Supplementary Figure 4.**
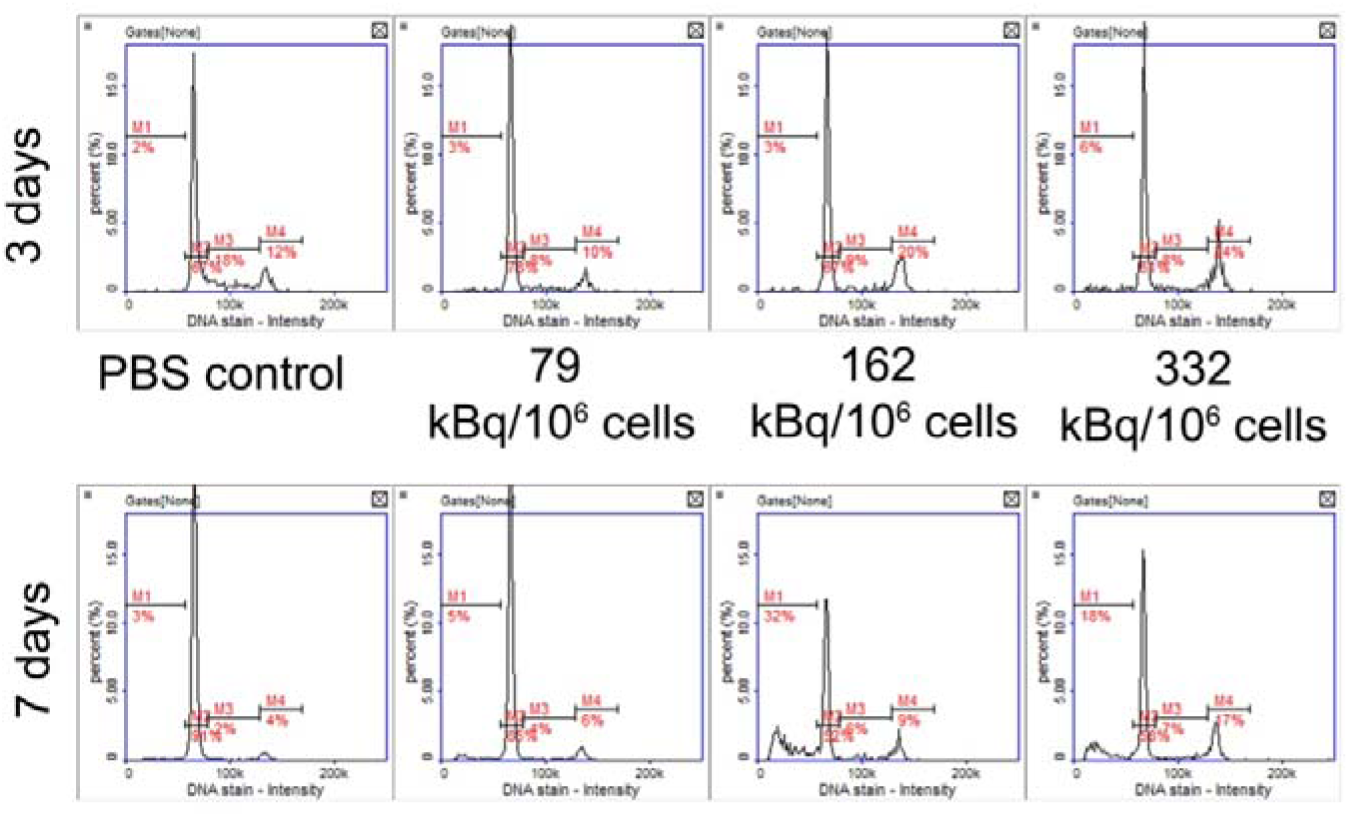
Cell cycle analysis was performed on populations of 10,000 cells per sample for the indicated doses and time points. At 3 days post dosing, the percentage of cells in S phase halved from 18% for the PBS control to 8% and 9% for the radiolabelled conditions. At the same time, the proportion of cells in G2 (the DNA damage checkpoint) doubled from 12% in the control condition to 20 and 24% in the top two radioactive doses. At 7 days post labelling, there was a large increase in the proportion of apoptotic cells in the top two radiolabelling doses, which was not seen at 3 days post labelling.

**Supplementary Figure 5.**
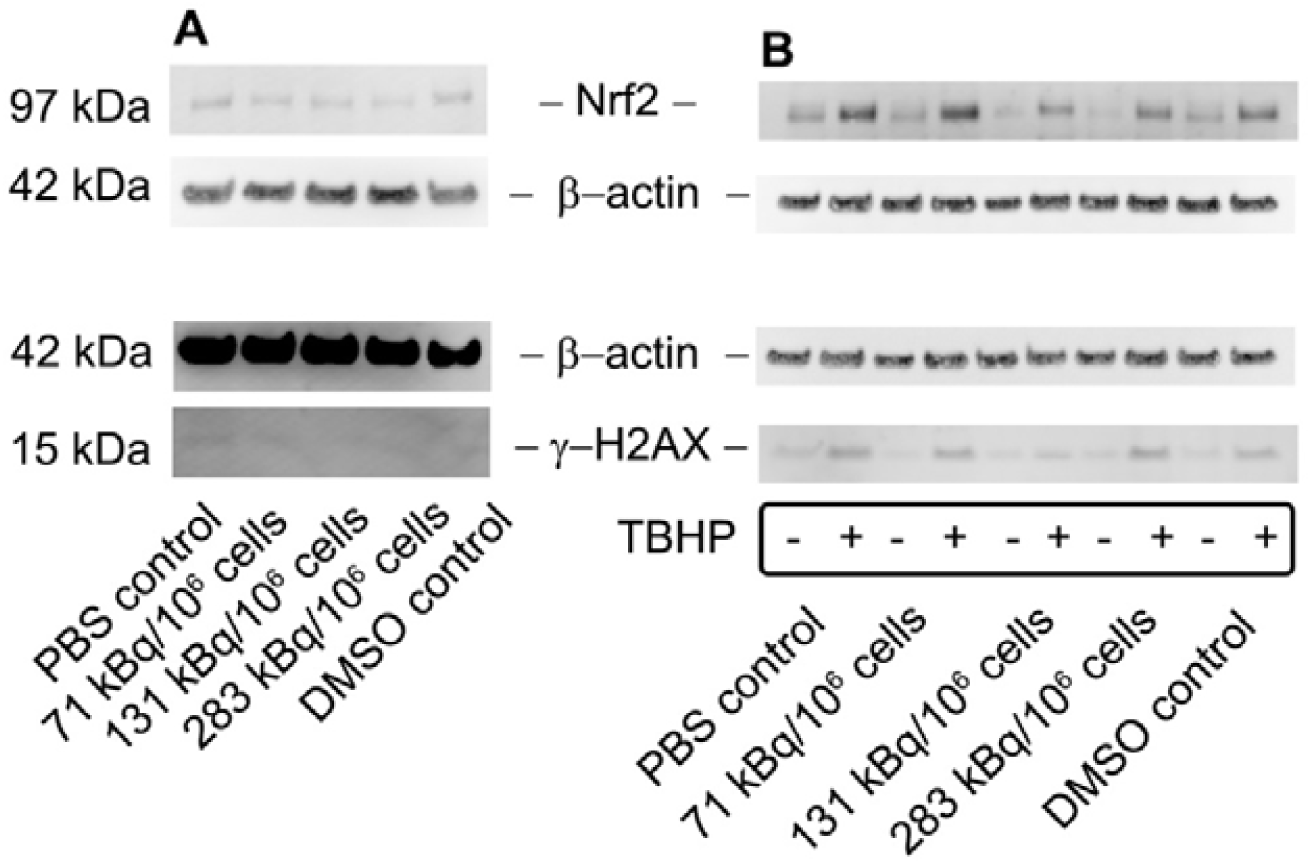
Western Blot analysis shows comparable expression of cell stress-associated protein Nrf2 and DNA damage-associated protein γ-H2AX in ^89^Zr-oxine labelled and unlabelled control cells at A. 7 days post radiolabelling. B. Treatment with 200 μM TBHP (tert-butyl-hydroperoxide) for 1 hour induces reactive oxygen stress and DNA damage, and upregulates Nrf2 and γ-H2AX expression in control and radiolabelled cell populations at 3 weeks post radiolabelling, confirming the sensitivity of these signalling pathways.

**Supplementary figure 6.**
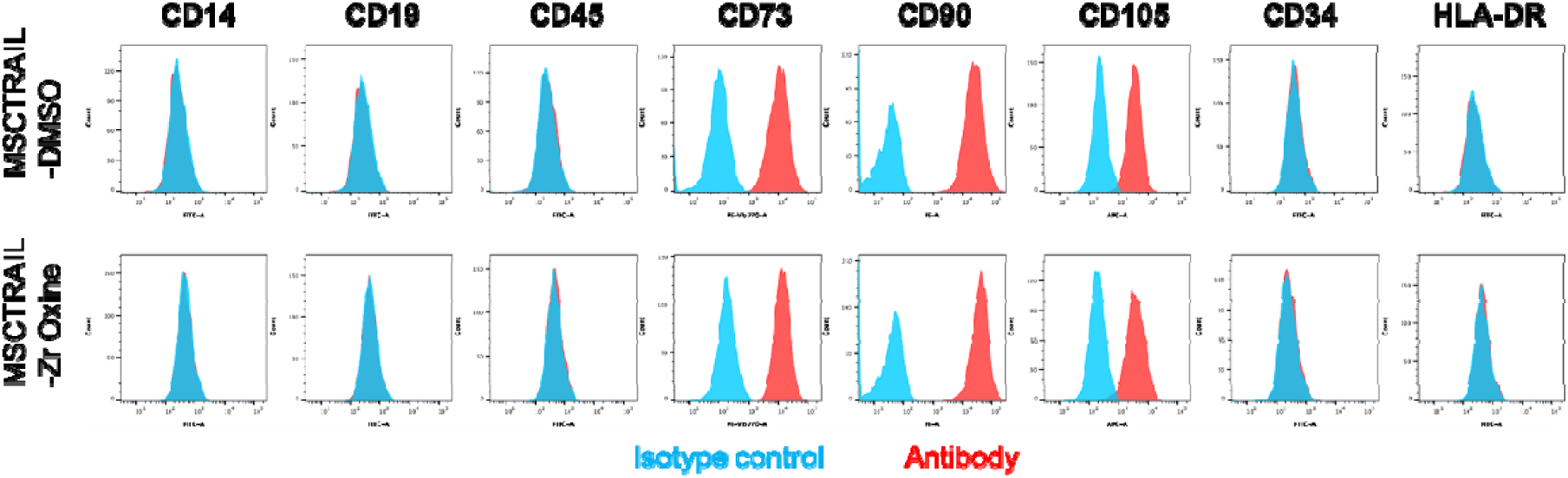
Cell surface marker phenotype was assessed using the ISCT-approved panel of MSC identification antibodies at 3 weeks post radiolabelling. Control (DMSO+PBS sham labelled) and radiolabelled (332 kBq/10^6^ cells) populations show positive staining for CD73, CD90, and CD 105, and negative staining for CD14, CD19, CD45, CD34 and HLA DR. PBS only (control) and lower radiolabelling doses (79, 162 kBq/10^6^ cells) gave comparable results (not shown).

**Supplementary figure 7.**
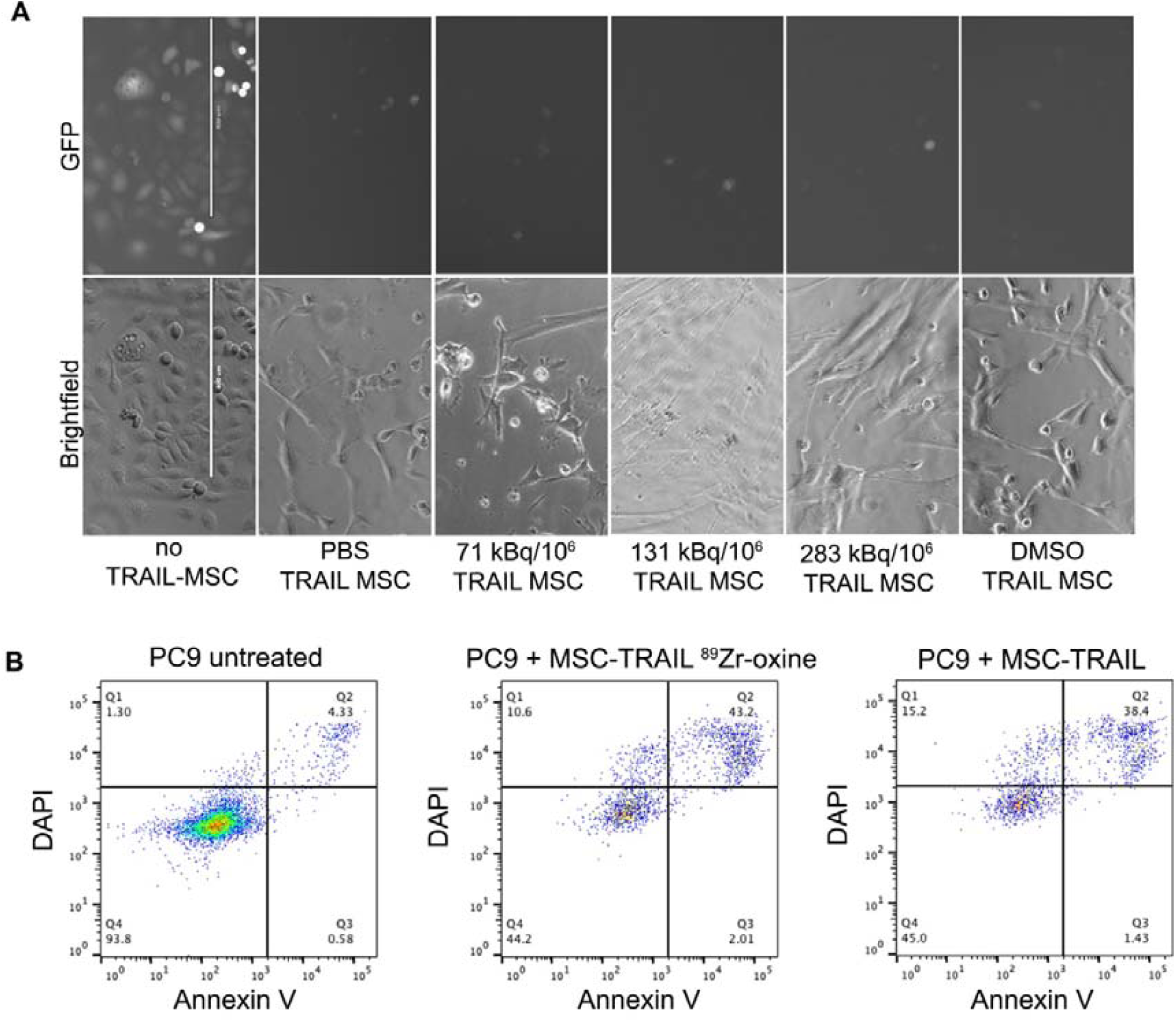
MSCTRAIL cells retain ability to induce apoptosis in cancer cells post radiolabelling. **A**. H28 Cells maintain viability in the absence of treatment (no MSCTRAIL), as seen by the presence of green fluorescent cells. Following treatment with sham dosed (PBS and DMSO MSCTRAIL), or radiolabelled MSCs at the indicated doses, GFP-expressing H28 cells are no longer seen and only MSCs are visible. Scale bars are 400 μm. **B**. Apoptosis is induced in PC9 cells following incubation with MSCTRAIL or radiolabelled MSCTRAIL (332 kBq/10^6^ cells).

**Supplementary figure 8.**
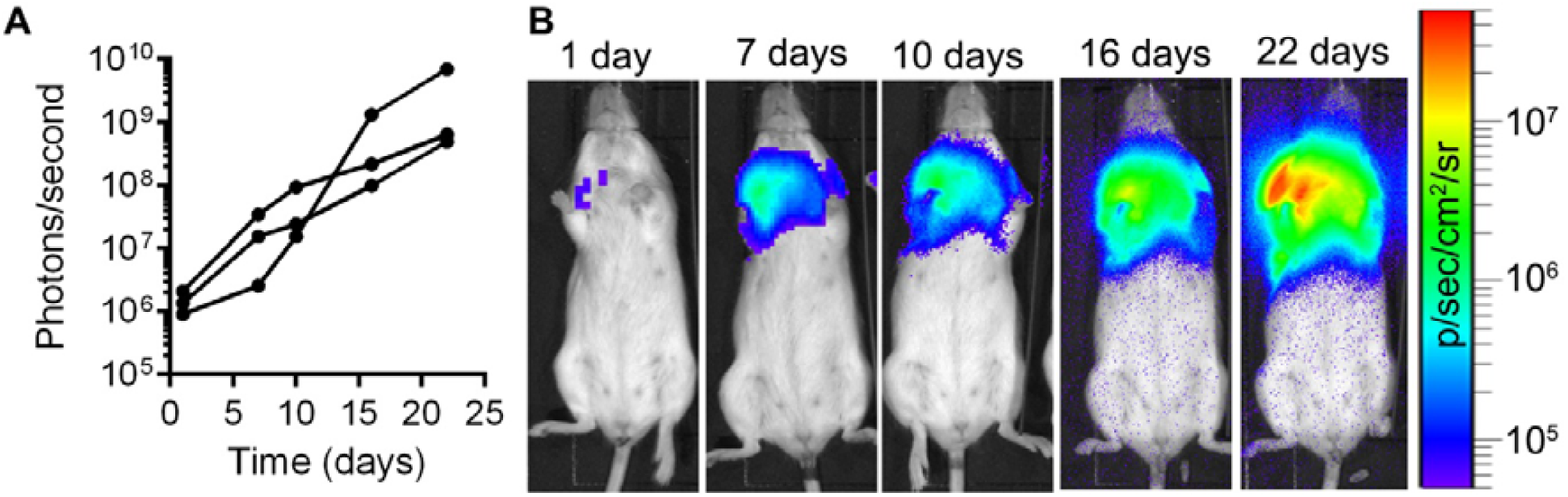
Growth of luciferase-expressing human lung mesothelioma cells (CRL2081) in mouse lungs was monitored using bioluminescence imaging at 20 minutes post injection of D-luciferin. **A**. Tumour growth represented as bioluminescent light output from a region of interest drawn over the lung area. Points show the light output from each time point post tumour implantation for three individual mice. **B**. Bioluminescence images from a representative mouse showing lung tumour growth up to 22 days post implantation. Light output intensity is indicated as colour heat-map overlaid on a bright-field image of the mouse.

**Supplementary figure 9.**
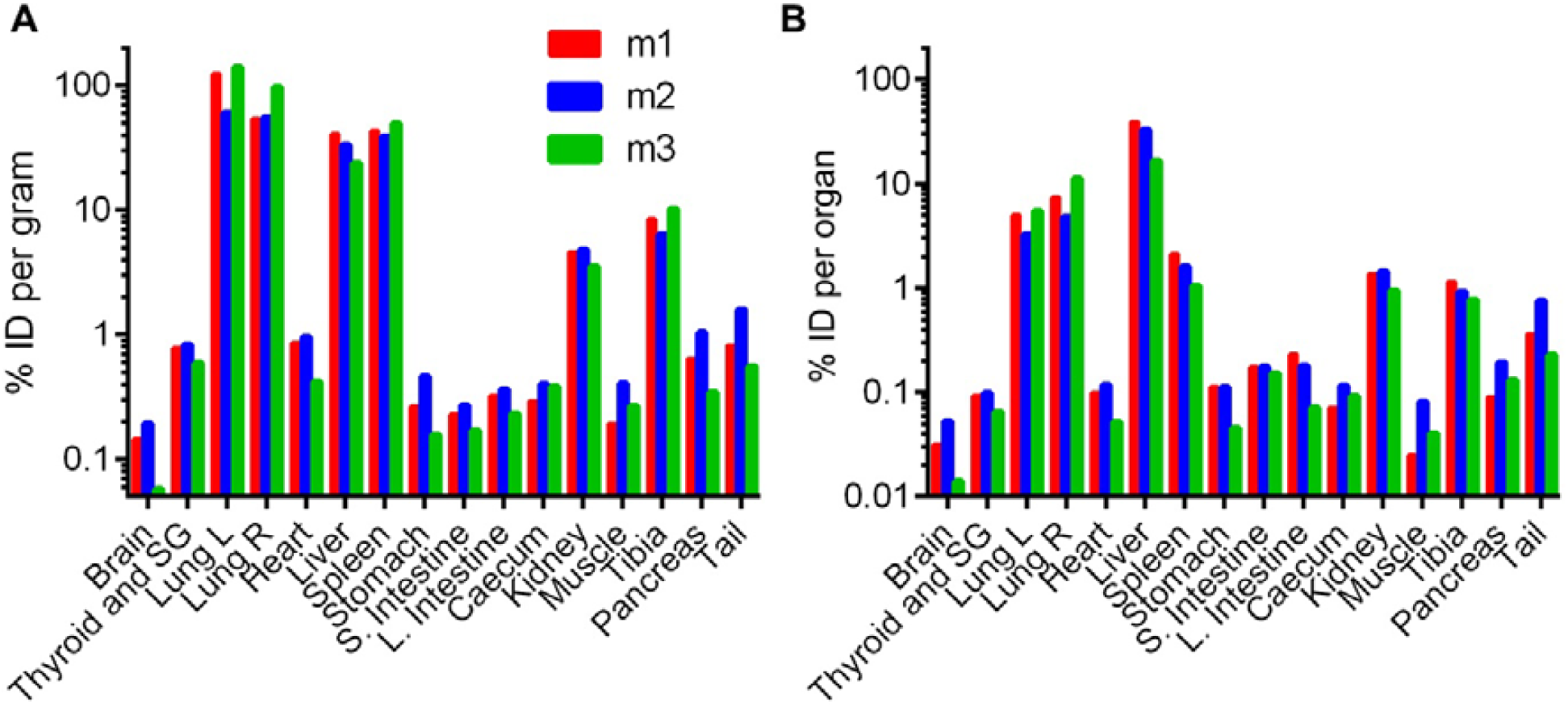
Biodistribution data at 10 days post ^89^Zr-oxine labelled MSCTRAIL injection showing decay-corrected ^89^Zr activity as % Injected dose A. per gram of wet tissue weight, and B. per organ for each of 3 mice.

**Supplementary figure 10.**
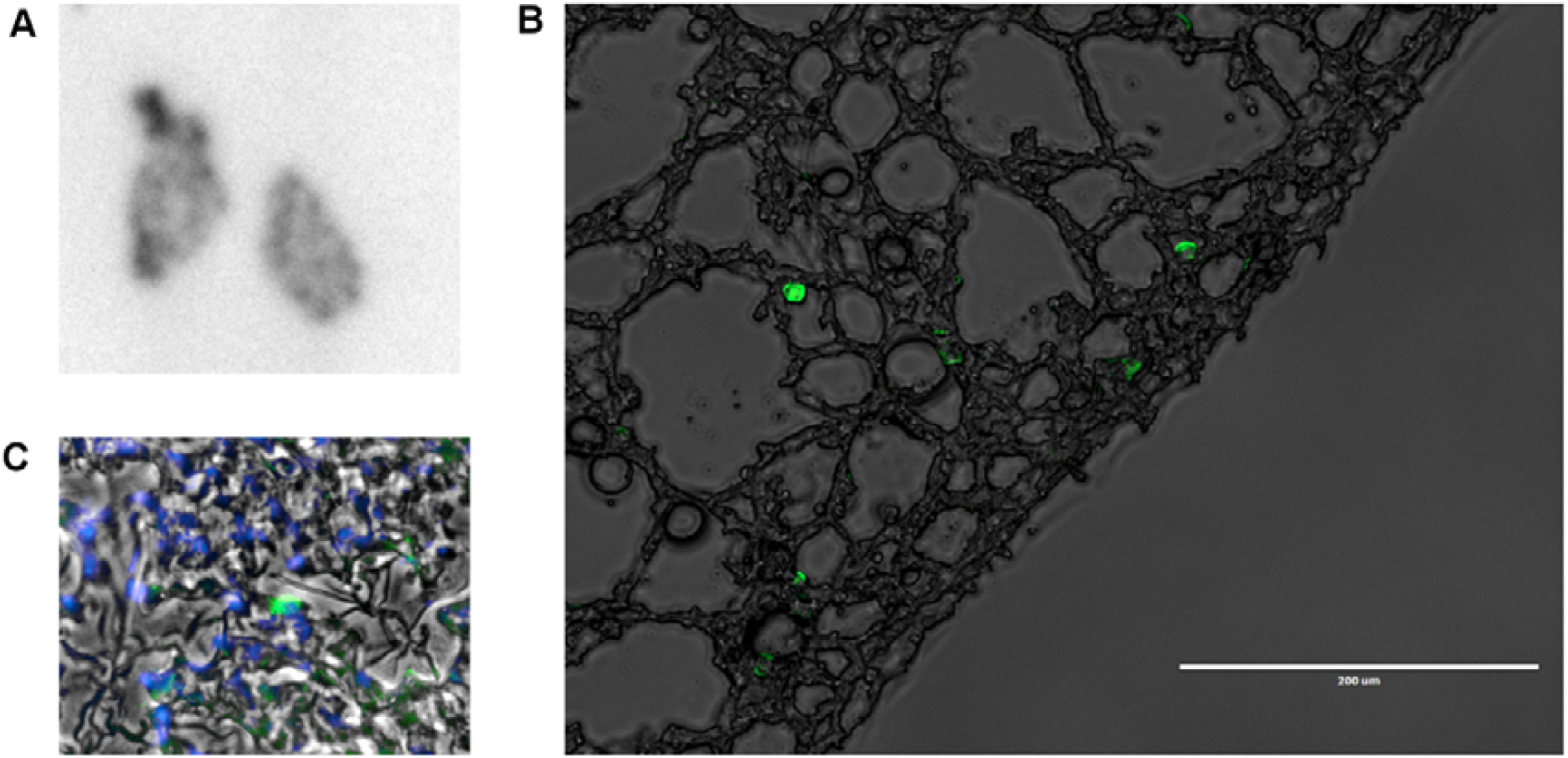
**A**. Autoradiography of cryosectioned (20 µm) lung tissue taken at 7 days post injection of ^89^Zr-oxine labelled cord-derived MSCs expressing ZsGreen and Luciferase B. Green fluorescence microscopy (ex. 470 nm; em. 510 nm) showing ZsGreen-expressing cord-derived MSCs overlaid onto phase contrast image of lung tissue shown in A. C. DAPI fluorescence showing nuclei, overlaid with green fluorescence showing ZsGreen-expressing cord-derived MSCs, overlaid onto phase contrast image of lung tissue shown in A.

**Supplementary Figure 11.**
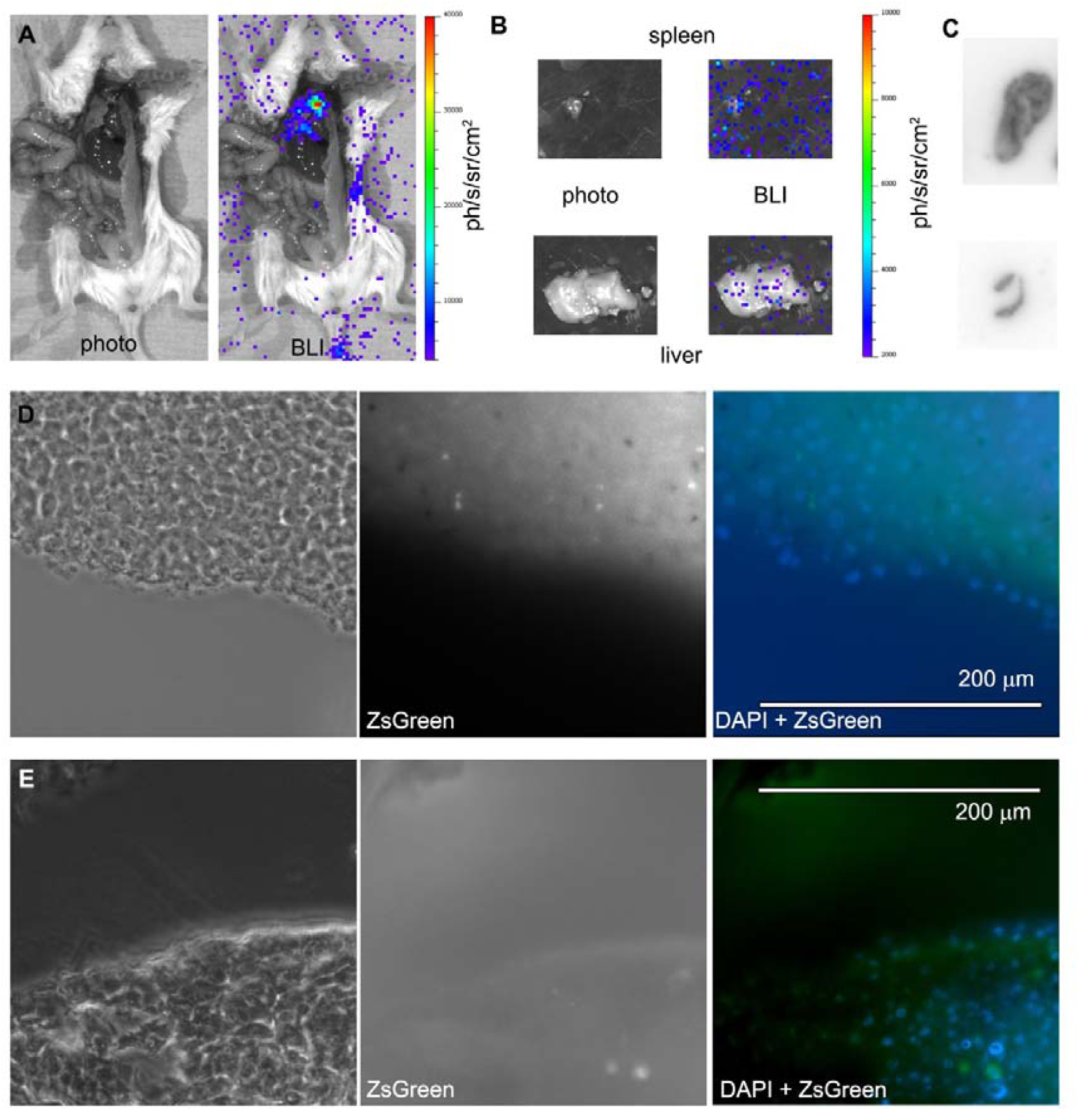
Ex vivo bioluminescence imaging showing **A**. opened mouse chest and abdominal cavity with signal visible in the lungs, and **B**. dissected liver and spleen showing background levels of signal. **C**. Autoradiography on cryosections of liver and spleen. Light and fluorescence microscopy showing slices of **D**. Liver, and E. spleen, showing nuclei stained with DAPI, and ZsGreen fluorescence showing possible debris from injected MSCs.

**Supplementary figure 12.**
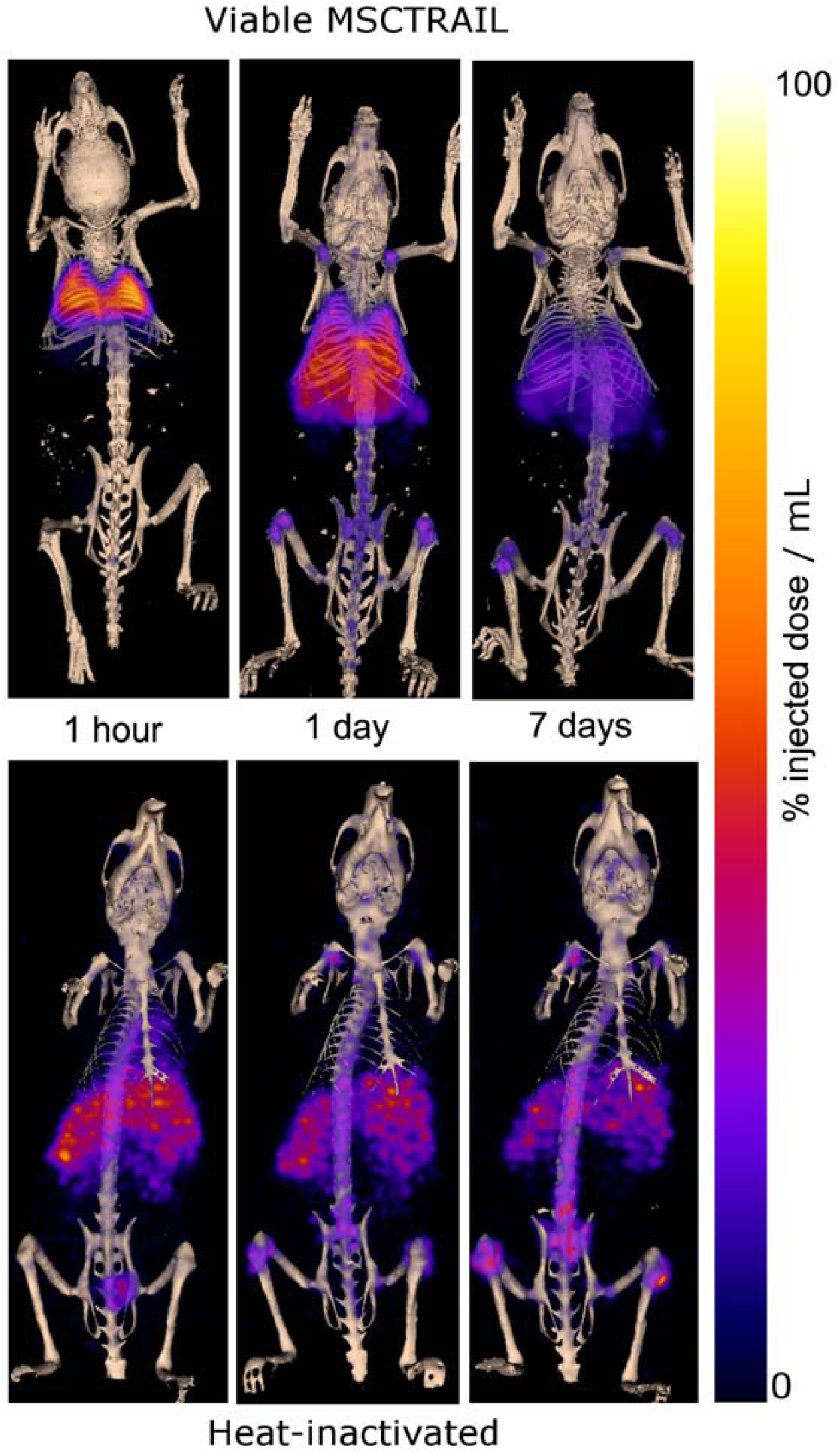
PET-CT maximum intensity projection images showing the distribution of signal at 1 hour, 1 day, and 7 days after intravenous injection of ^89^Zr-oxine labelled MSCTRAIL and ^89^Zr-oxine labelled cells that have been heat-inactivated (90°C for 20 minutes).

**Table S1.**
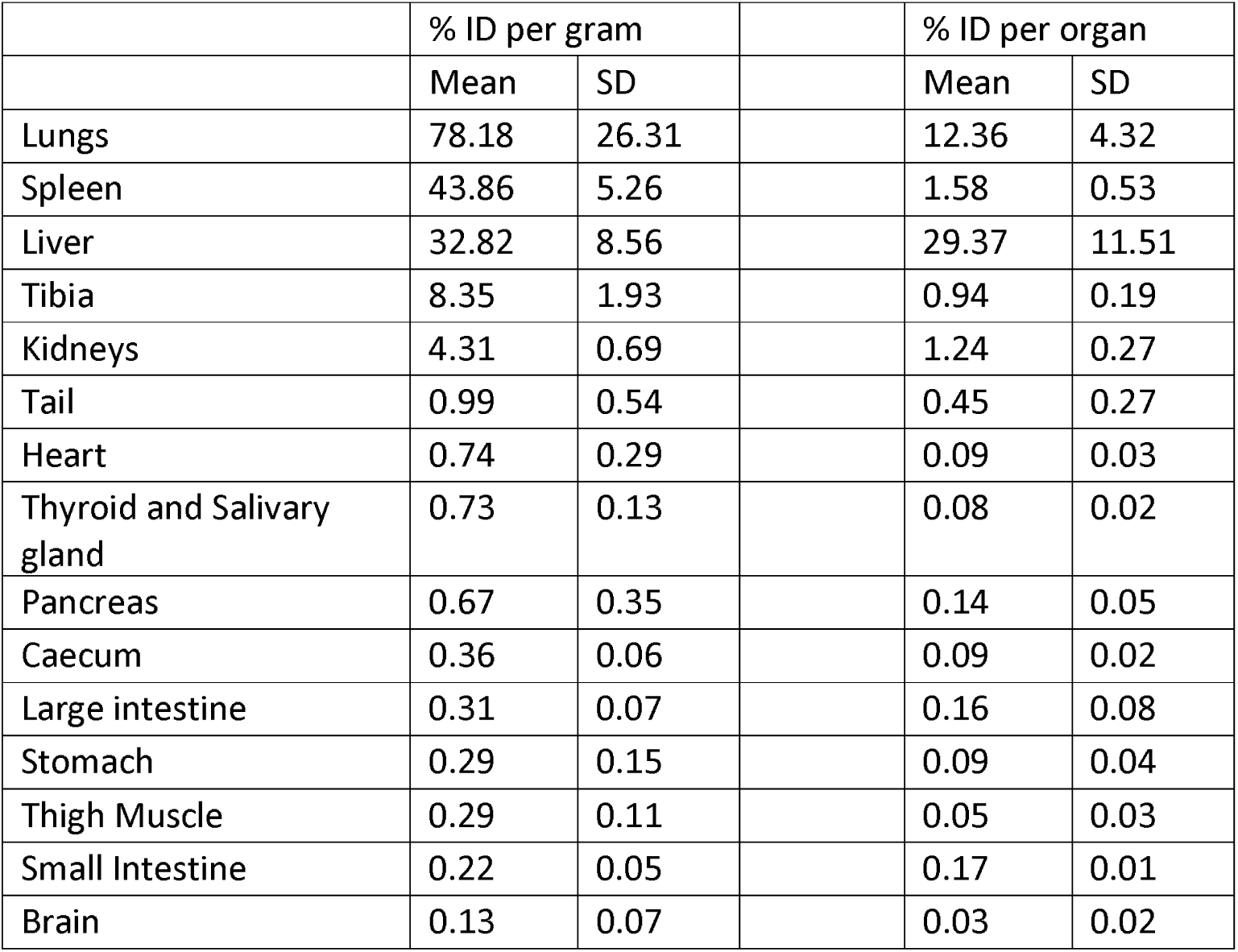
*Ex vivo* biodistribution data showing ^89^Zr activity measurements in the indicated organs at 10 days post injection of ^89^Zr-oxine labelled MSCTRAIL. Organs are ordered in descending values of % injected dose per gram tissue. N=3 animals and SD indicates standard deviation.

|  | % ID per gram |  |  | % ID per organ |  |
| --- | --- | --- | --- | --- | --- |
|  | Mean | SD |  | Mean | SD |
| Lungs | 78.18 | 26.31 |  | 12.36 | 4.32 |
| Spleen | 43.86 | 5.26 |  | 1.58 | 0.53 |
| Liver | 32.82 | 8.56 |  | 29.37 | 11.51 |
| Tibia | 8.35 | 1.93 |  | 0.94 | 0.19 |
| Kidneys | 4.31 | 0.69 |  | 1.24 | 0.27 |
| Tail | 0.99 | 0.54 |  | 0.45 | 0.27 |
| Heart | 0.74 | 0.29 |  | 0.09 | 0.03 |
| Thyroid and Salivary gland | 0.73 | 0.13 |  | 0.08 | 0.02 |
| Pancreas | 0.67 | 0.35 |  | 0.14 | 0.05 |
| Caecum | 0.36 | 0.06 |  | 0.09 | 0.02 |
| Large intestine | 0.31 | 0.07 |  | 0.16 | 0.08 |
| Stomach | 0.29 | 0.15 |  | 0.09 | 0.04 |
| Thigh Muscle | 0.29 | 0.11 |  | 0.05 | 0.03 |
| Small Intestine | 0.22 | 0.05 |  | 0.17 | 0.01 |
| Brain | 0.13 | 0.07 |  | 0.03 | 0.02 |

**Table S2.** Equivalent doses for a range of organs/tissues calculated using OLINDA in a male phantom model. The dose is quoted in in mSv per MBq injected, and also in mSv for 37 and 100 MBq injected activities.

| **Organ/Tissue** | **Equivalent Dose per MBq injected (mSv/MBq) for Adult Male** | **Equivalent Dose for 37 MBq injected activity (mSv) for Adult Male** | **Equivalent Dose for 100 MBq injected activity (mSv) for Adult Male** |
| --- | --- | --- | --- |
| Adrenals | 0.68 (0.56-0.87) | 25.0 | 67.6 |
| Brain | 0.07 (0.05-0.10) | 2.6 | 7.2 |
| Gallbladder Wall | 0.57 (0.46-0.67) | 21.2 | 57.4 |
| LLI Wall | 0.06 (0.04-0.07) | 2.1 | 5.6 |
| Small Intestine | 0.13 (0.10-0.16) | 4.8 | 13.1 |
| Stomach Wall | 0.35 (0.28-0.47) | 12.8 | 34.6 |
| ULI Wall | 0.17 (0.14-0.21) | 6.3 | 17.1 |
| Heart Wall | 0.80 (0.66-1.06) | 29.6 | 80.0 |
| Kidneys | 0.61 (0.48-0.73) | 22.6 | 61.0 |
| Liver | 1.86 (1.49-2.14) | 68.8 | 186.0 |
| Lungs | 5.09 (3.97-6.89) | 188.2 | 508.7 |
| Muscle | 0.23 (0.19-0.31) | 8.7 | 23.4 |
| Pancreas | 0.58 (0.46-0.76) | 21.3 | 57.5 |
| Red Marrow | 0.33 (0.27-0.44) | 12.1 | 32.7 |
| Osteogenic Cells | 0.55 (0.42-0.75) | 20.5 | 55.3 |
| Skin | 0.13 (0.11-0.18) | 4.9 | 13.3 |
| Spleen | 2.12 (1.49-3.36) | 78.3 | 211.7 |
| Testes | 0.02 (0.02-0.03) | 0.8 | 2.3 |
| Thymus | 0.51 (0.41-0.69) | 18.9 | 51.0 |
| Thyroid | 0.19 (0.15-0.25) | 6.9 | 18.6 |
| Urinary Bladder Wall | 0.04 (0.03-0.05) | 1.3 | 3.5 |

**Table S3.** Equivalent doses for a range of organs/tissues calculated using OLINDA in a female phantom model. The dose is quoted in in mSv per MBq injected, and also in mSv for 37 and 100 MBq injected activities. Table 2. Organ doses for adult female phantom model

| Organ/Tissue | Equivalent Dose per MBq injected (mSv/MBq) for Adult Female | Equivalent Dose for 37 MBq injected activity (mSv) for Adult Female | Equivalent Dose for 100 MBq injected activity (mSv) for Adult Female |
| --- | --- | --- | --- |
| Adrenals | 0.86 (0.70-1.11) | 31.8 | 85.9 |
| Brain | 0.08 (0.06-0.11) | 3.0 | 8.1 |
| Breasts | 0.53 (0.43-0.71) | 19.5 | 52.8 |
| Gallbladder Wall | 0.70 (0.56-0.82) | 25.8 | 69.6 |
| LLI Wall | 0.08 (0.06-0.10) | 2.8 | 7.6 |
| Small Intestine | 0.17 (0.13-0.21) | 6.2 | 16.8 |
| Stomach Wall | 0.46 (0.37-0.61) | 16.9 | 45.8 |
| ULI Wall | 0.21 (0.17-0.26) | 7.9 | 21.2 |
| Heart Wall | 1.05 (0.87-1.39) | 38.8 | 104.9 |
| Kidneys | 0.72 (0.56-0.86) | 26.5 | 71.7 |
| Liver | 2.39 (1.92-2.75) | 88.4 | 239.0 |
| Lungs | 6.58 (5.13-8.91) | 243.3 | 657.7 |
| Muscle | 0.30 (0.25-0.4) | 11.2 | 30.3 |
| Ovaries | 0.09 (0.07-0.12) | 3.3 | 9.0 |
| Pancreas | 0.76 (0.61-1.01) | 28.2 | 76.3 |
| Red Marrow | 0.38 (0.31-0.51) | 14.1 | 38.1 |
| Osteogenic Cells | 0.71 (0.54-0.97) | 26.3 | 71.2 |
| Skin | 0.16 (0.13-0.21) | 5.9 | 16.0 |
| Spleen | 2.57 (1.82-4.07) | 95.2 | 257.3 |
| Thymus | 0.61 (0.49-0.82) | 22.6 | 61.2 |
| Thyroid | 0.23 (0.19-0.32) | 8.6 | 23.4 |
| Urinary Bladder Wall | 0.05 (0.04-0.06) | 1.8 | 5.0 |
| Uterus | 0.08 (0.06-0.10) | 3.0 | 8.2 |

**Table S4.** Comparison of estimated human dosing between ^89^ Zr-oxine cell injection and published studies using ^89^ Zr-antibodies.

| Source | Zr-89 Oxine TRAIL MSCs (37 MBq, Male) | Zr-89 Oxine TRAIL MSCs (100 MBq, Male) | Zr-89 Oxine TRAIL MSCs (37 MBq, Female) | Zr-89 Oxine TRAIL MSCs (100 MBq, Female) | Zr-89 Chimeric Monoclonal Antibody U36 (75 MBq, Female) [2] | Zr-89 Ibritumomab Tiuxetan (70 MBq) [3] |
| --- | --- | --- | --- | --- | --- | --- |
| Effective Dose per MBq injected (mSv/MBq) | 0.87 (0.72-1.16) |  | 1.12 (0.92-1.49) |  | 0.66 | 0.55 |
| Effective Dose (mSv) | 32.2 | 87.1 | 41.4 | 111.8 | 49.5 | 38.5 |
| Highest organ equivalent dose (mSv/MBq) | 5.09 (3.97-6.89) Lungs |  | 6.58 (5.13-8.91) Lungs |  | 1.35 (Liver) | 1.36 (Liver) |
| Highest organ equivalent dose (mSv) | 188.2 | 508.7 | 243.3 | 657.7 | 101.3 | 95.2 |

## Zr-89 Oxine MSC-TRAIL: Human Dosimetry Estimates based upon Pre-clinical Data

Pre-clinical data from the Centre for Advanced Biomedical Imaging (CABI) at UCL was used to estimate organ doses and an effective dose in adult humans (male and female) for the administration of ^89^Zr-Oxine labelled MSC-TRAIL.

The estimated effective dose is 0.87 mSv/MBq (range 0.72-1.16, n=3) for males, and 1.12 mSv/MBq (range 0.92-1.49, n=3) for females. For an administered activity of 37 MBq the effective dose is estimated as 32.2 (26.4-42.9) mSv for males, and 41.1 (34.0-55.1) mSv for females. For an administered activity of 100 MBq the effective dose is estimated as 87.1 (71.6-116.0) mSv for males, and 111.8 (91.9-149.0) mSv for females.

These are high doses in comparison with routine clinical PET imaging, for example an FDG PET is approximately 7.6 mSv for a 400 MBq administration [1]. The doses are more comparable with those reported for other Zr-89 labelled PET studies, but are still approximately 30-100% higher for equivalent administered activities [2] [3]. The highest organ equivalent dose is to the lungs, and is estimated to be 5.1 mSv/MBq (range 4.0-6.9) for males and 5.82 mSv/MBq (range 6.6-8.9) for females.

### Pre-clinical Data

The pre-clinical data included the % of injected dose per gram of tissue (%ID/g) in mice for the Lungs, Liver, Spleen, Kidneys, and Bone, derived from PET/CT imaging at 1, 24, 48, and 168 hours post administration in three mice (for one mouse the 1 hour time point was not available). At 240 hours post administration the %ID/g was measured ex vivo for the following organs; Brain, Thyroid, Lungs, Heart, Liver, Spleen, Stomach, Small Intestine, Large Intestine, Caecum, Kidney, Muscle, Tibia, Pancreas, and Tail. Organ masses were also measured. The whole body weight of each mouse was estimated as 20 g.

For the PET/CT derived data, the decay corrected %ID/g for each organ (Lungs, Liver, Spleen, Kidneys, and Bone) was calculated for each mouse. The ex vivo data at 240 hours was not used for these organs, as the different method of measurement lead to a poor curve fit. The organs with PET/CT data accounted for 89-91% of the injected activity at 1 hour.

### Human Dosimetry Estimate

The mouse data was extrapolated to human data using equations provided on p83 and 84 of Stabin, *Fundamentals of Nuclear Medicine Dosimetry* [4]. This included a conversion from mouse %ID/g to human %ID/organ using the measured mouse organ and whole-body masses and human phantom model masses from OLINDA, and a transformation of the timescale to account for differences in metabolic rate for species of different body mass [4]. The %ID/organ was normalised so that the total summed across the lungs, liver, kidneys, spleen, and bone was equal to 100%, as no rapid excretion is expected.

Curve fitting of time-activity plots was performed either in OLINDA (version 1.0, 2003) [5] or MATLAB, using either a multi-phase exponential decay model or a bi-exponential decay model that includes an uptake phase. The MIRD dosimetry methodology was applied to obtain the residence time for each organ.

The residence time was inputted into OLINDA. Organ doses were calculated using the adult male and adult female models. To match the OLINDA kinetics input form, all bone activity was assumed to be in the cortical bone.

## References

1. Siegel RL, Miller KD, Jemal A. Cancer statistics, 2018. CA: a cancer journal for clinicians. 2018;68:7–30. doi:10.3322/caac.21442.

2. Fischbach MA, Bluestone JA, Lim WA. Cell-based therapeutics: the next pillar of medicine. Science translational medicine. 2013;5:179ps7. doi:10.1126/scitranslmed.3005568.

3. Bulte JWM, Daldrup-Link HE. Clinical Tracking of Cell Transfer and Cell Transplantation: Trials and Tribulations. Radiology. 2018;289:604–15. doi:10.1148/radiol.2018180449.

4. Wang J, Jokerst JV. Stem Cell Imaging: Tools to Improve Cell Delivery and Viability. Stem cells international. 2016;2016:9240652. doi:10.1155/2016/9240652.

5. Scarfe L, Brillant N, Kumar JD, Ali N, Alrumayh A, Amali M, et al. Preclinical imaging methods for assessing the safety and efficacy of regenerative medicine therapies. NPJ Regenerative medicine. 2017;2:28. doi:10.1038/s41536-017-0029-9.

6. Thakrar RM, Sage EK, Janes SM. Combined cell-gene therapy for lung cancer: rationale, challenges and prospects. Expert opinion on biological therapy. 2016;16:853–7. doi:10.1080/14712598.2016.1188074.

7. Sage EK, Thakrar RM, Janes SM. Genetically modified mesenchymal stromal cells in cancer therapy. Cytotherapy. 2016;18:1435–45. doi:10.1016/j.jcyt.2016.09.003.

8. Kolluri KK, Laurent GJ, Janes SM. Mesenchymal stem cells as vectors for lung cancer therapy. Respiration; international review of thoracic diseases. 2013;85:443–51. doi:10.1159/000351284.

9. Lourenco S, Teixeira VH, Kalber T, Jose RJ, Floto RA, Janes SM. Macrophage migration inhibitory factor-CXCR4 is the dominant chemotactic axis in human mesenchymal stem cell recruitment to tumors. Journal of immunology. 2015;194:3463–74. doi:10.4049/jimmunol.1402097.10.

10. Loebinger MR, Kyrtatos PG, Turmaine M, Price AN, Pankhurst Q, Lythgoe MF, et al. Magnetic resonance imaging of mesenchymal stem cells homing to pulmonary metastases using biocompatible magnetic nanoparticles. Cancer research. 2009;69:8862–7. doi:10.1158/0008-5472.CAN-09-1912.

11. Ferris TJ, Charoenphun P, Meszaros LK, Mullen GE, Blower PJ, Went MJ. Synthesis and characterisation of zirconium complexes for cell tracking with Zr-89 by positron emission tomography. Dalton transactions. 2014;43:14851–7. doi:10.1039/c4dt01928h.

12. Charoenphun P, Meszaros LK, Chuamsaamarkkee K, Sharif-Paghaleh E, Ballinger JR, Ferris TJ, et al. [(89)Zr]oxinate4 for long-term in vivo cell tracking by positron emission tomography. European journal of nuclear medicine and molecular imaging. 2015;42:278–87. doi:10.1007/s00259-014-2945-x.

13. Sato N, Wu H, Asiedu KO, Szajek LP, Griffiths GL, Choyke PL. (89)Zr-Oxine Complex PET Cell Imaging in Monitoring Cell-based Therapies. Radiology. 2015;275:490–500. doi:10.1148/radiol.15142849.

14. Rahmim A, Zaidi H. PET versus SPECT: strengths, limitations and challenges. Nuclear medicine communications. 2008;29:193–207. doi:10.1097/MNM.0b013e3282f3a515.

15. Cherry SR, Jones T, Karp JS, Qi J, Moses WW, Badawi RD. Total-Body PET: Maximizing Sensitivity to Create New Opportunities for Clinical Research and Patient Care. Journal of nuclear medicine: official publication, Society of Nuclear Medicine. 2018;59:3–12. doi:10.2967/jnumed.116.184028.

16. Weist MR, Starr R, Aguilar B, Chea J, Miles JK, Poku E, et al. PET of Adoptively Transferred Chimeric Antigen Receptor T Cells with (89)Zr-Oxine. Journal of nuclear medicine: official publication, Society of Nuclear Medicine. 2018;59:1531–7. doi:10.2967/jnumed.117.206714.

17. Man F, Lim L, Volpe A, Gabizon A, Shmeeda H, Draper B, et al. In Vivo PET Tracking of (89)Zr-Labeled Vgamma9Vdelta2 T Cells to Mouse Xenograft Breast Tumors Activated with Liposomal Alendronate. Molecular therapy: the journal of the American Society of Gene Therapy. 2019;27:219–29. doi:10.1016/j.ymthe.2018.10.006.

18. Asiedu KO, Ferdousi M, Ton PT, Adler SS, Choyke PL, Sato N. Bone marrow cell homing to sites of acute tibial fracture: (89)Zr-oxine cell labeling with positron emission tomographic imaging in a mouse model. EJNMMI research. 2018;8:109. doi:10.1186/s13550-018-0463-8.

19. Asiedu KO, Koyasu S, Szajek LP, Choyke PL, Sato N. Bone Marrow Cell Trafficking Analyzed by (89)Zr-oxine Positron Emission Tomography in a Murine Transplantation Model. Clinical cancer research: an official journal of the American Association for Cancer Research. 2017;23:2759–68. doi:10.1158/1078-0432.CCR-16-1561.

20. Yuan Z, Lourenco Sda S, Sage EK, Kolluri KK, Lowdell MW, Janes SM. Cryopreservation of human mesenchymal stromal cells expressing TRAIL for human anti-cancer therapy. Cytotherapy. 2016;18:860–9. doi:10.1016/j.jcyt.2016.04.005.

21. Dominici M, Le Blanc K, Mueller I, Slaper-Cortenbach I, Marini F, Krause D, et al. Minimal criteria for defining multipotent mesenchymal stromal cells. The International Society for Cellular Therapy position statement. Cytotherapy. 2006;8:315–7. doi:10.1080/14653240600855905.

22. Stabin MG, Sparks RB, Crowe E. OLINDA/EXM: the second-generation personal computer software for internal dose assessment in nuclear medicine. Journal of nuclear medicine: official publication, Society of Nuclear Medicine. 2005;46:1023–7.

23. Stabin MG. Fundamentals of Nuclear Medicine Dosimetry: Springer; 2008.

24. Hofmann M, Wollert KC, Meyer GP, Menke A, Arseniev L, Hertenstein B, et al. Monitoring of bone marrow cell homing into the infarcted human myocardium. Circulation. 2005;111:2198–202. doi:10.1161/01.CIR.0000163546.27639.AA.

25. Uribe-Herranz M, Bittinger K, Rafail S, Guedan S, Pierini S, Tanes C, et al. Gut microbiota modulates adoptive cell therapy via CD8alpha dendritic cells and IL-12. JCI Insight. 2018;3. doi:10.1172/jci.insight.94952.

26. Dimmeler S, Leri A. Aging and disease as modifiers of efficacy of cell therapy. Circulation research. 2008;102:1319–30. doi:10.1161/CIRCRESAHA.108.175943.

27. Marks PW, Witten CM, Califf RM. Clarifying Stem-Cell Therapy’s Benefits and Risks. The New England journal of medicine. 2017;376:1007–9. doi:10.1056/NEJMp1613723.

28. Dijkers EC, Oude Munnink TH, Kosterink JG, Brouwers AH, Jager PL, de Jong JR, et al. Biodistribution of 89Zr-trastuzumab and PET imaging of HER2-positive lesions in patients with metastatic breast cancer. Clin Pharmacol Ther. 2010;87:586–92. doi:10.1038/clpt.2010.12.

29. Gaykema SB, Brouwers AH, Lub-de Hooge MN, Pleijhuis RG, Timmer-Bosscha H, Pot L, et al. 89Zr-bevacizumab PET imaging in primary breast cancer. Journal of nuclear medicine: official publication, Society of Nuclear Medicine. 2013;54:1014–8. doi:10.2967/jnumed.112.117218.

30. Borjesson PK, Jauw YW, Boellaard R, de Bree R, Comans EF, Roos JC, et al. Performance of immuno-positron emission tomography with zirconium-89-labeled chimeric monoclonal antibody U36 in the detection of lymph node metastases in head and neck cancer patients. Clinical cancer research: an official journal of the American Association for Cancer Research. 2006;12:2133–40. doi:10.1158/1078-0432.CCR-05-2137.

31. Rizvi SNF, Visser OJ, Vosjan M, van Lingen A, Hoekstra OS, Zijlstra JM, et al. Biodistribution, radiation dosimetry and scouting of (90)Y-ibritumomab tiuxetan therapy in patients with relapsed B-cell non-Hodgkin’s lymphoma using (89)Zr-ibritumomab tiuxetan and PET. European journal of nuclear medicine and molecular imaging. 2012;39:512–20.

32. Mendicino M, Bailey AM, Wonnacott K, Puri RK, Bauer SR. MSC-based product characterization for clinical trials: an FDA perspective. Cell stem cell. 2014;14:141–5. doi:10.1016/j.stem.2014.01.013.

33. Heathman TR, Nienow AW, McCall MJ, Coopman K, Kara B, Hewitt CJ. The translation of cell-based therapies: clinical landscape and manufacturing challenges. Regen Med. 2015;10:49–64. doi:10.2217/rme.14.73.

34. Fan CG, Zhang QJ, Zhou JR. Therapeutic potentials of mesenchymal stem cells derived from human umbilical cord. Stem cell reviews. 2011;7:195–207. doi:10.1007/s12015-010-9168-8.

35. Nagamura-Inoue T, He H. Umbilical cord-derived mesenchymal stem cells: Their advantages and potential clinical utility. World journal of stem cells. 2014;6:195–202. doi:10.4252/wjsc.v6.i2.195.

36. Moroz MA, Zhang H, Lee J, Moroz E, Zurita J, Shenker L, et al. Comparative Analysis of T Cell Imaging with Human Nuclear Reporter Genes. Journal of nuclear medicine: official publication, Society of Nuclear Medicine. 2015;56:1055–60. doi:10.2967/jnumed.115.159855.

37. Abou DS, Ku T, Smith-Jones PM. In vivo biodistribution and accumulation of 89Zr in mice. Nuclear medicine and biology. 2011;38:675–81. doi:10.1016/j.nucmedbio.2010.12.011.

38. Patrick PS, Hammersley J, Loizou L, Kettunen MI, Rodrigues TB, Hu DE, et al. Dual-modality gene reporter for in vivo imaging. Proceedings of the National Academy of Sciences of the United States of America. 2014;111:415–20. doi:10.1073/pnas.1319000111.

39. Yuan Z, Kolluri KK, Sage EK, Gowers KH, Janes SM. Mesenchymal stromal cell delivery of full-length tumor necrosis factor-related apoptosis-inducing ligand is superior to soluble type for cancer therapy. Cytotherapy. 2015;17:885–96. doi:10.1016/j.jcyt.2015.03.603.

## References

1. Comittee A. Arsac notes for guidance notes for guidance on the clinical administration of radiopharmaceuticals and use of sealed radioactive sources. (2016).

2. Borjesson PK, Jauw YW, Boellaard R et al. Performance of immuno-positron emission tomography with zirconium-89-labeled chimeric monoclonal antibody u36 in the detection of lymph node metastases in head and neck cancer patients. Clinical cancer research: an official journal of the American Association for Cancer Research 12(7 Pt 1), 2133–2140 (2006).

3. Rizvi SNF, Visser OJ, Vosjan M et al. Biodistribution, radiation dosimetry and scouting of (90)y-ibritumomab tiuxetan therapy in patients with relapsed b-cell non-hodgkin’s lymphoma using (89)zr-ibritumomab tiuxetan and pet. European journal of nuclear medicine and molecular imaging 39(3), 512–520 (2012).

4. Stabin MG. Fundamentals of nuclear medicine dosimetry. Springer, (2008).

5. Stabin MG, Sparks RB, Crowe E. Olinda/exm: The second-generation personal computer software for internal dose assessment in nuclear medicine. Journal of nuclear medicine: official publication, Society of Nuclear Medicine 46(6), 1023–1027 (2005).

